# Transcriptional repression of central spindle factors controls endomitosis in the *C. elegans* intestine

**DOI:** 10.1101/2025.08.04.668394

**Authors:** Ramon Barrull-Mascaró, Sonia Veltkamp, Samia Boutaibi, Lotte M. van Rijnberk, Rebecca Lippmann, Matilde Galli

## Abstract

During development, many cell types transition from canonical to non-canonical cell cycles, such as endomitosis and endoreplication, in which they duplicate their DNA but do not divide, giving rise to polyploidy. Little is known on the regulation of endomitosis, where cells enter M phase, but do not perform cytokinesis. Here, we investigate how cells initiate and execute endomitosis in the *C. elegans* intestine and find that endomitotic cells fail to assemble a central spindle or initiate cytokinetic furrowing. We find that endomitotic cells transcriptionally repress multiple cytokinesis regulators, especially the central spindle factors ZEN-4^Mklp^^1^, CYK-4^RacGap^^1^ and SPD-1^Prc^^1^. Intestinal cells lose the capacity to perform cytokinesis in late embryogenesis, and we find that the conserved DREAM (DP, RB-related, E2F and MuvB) complex is involved in the repression of central spindle genes. Together, our work demonstrates that the transition to endomitosis is a well-defined switch that relies on the transcriptional repression of cytokinesis genes.

## INTRODUCTION

The canonical cell cycle consists of phases of growth (gap or G phase) and DNA synthesis (S phase), and culminates with the generation of two daughter cells during M phase, when cells undergo nuclear division (mitosis) and cellular division (cytokinesis)^1^. During development of multicellular organisms, different cell types alter their cell cycles to generate organs and tissues with specific sizes, shapes and functions^2–4^. For example, cells can modulate their cycling speed or the stringency of their cell-cycle checkpoints, or they can permanently arrest or exit the cell cycle^5^. In addition, many cell types transition to non-canonical cell cycles, in which they duplicate their DNA but do not generate two daughter cells, instead giving rise to polyploid cells that have extra copies of the genomic DNA^6^.

Two types of non-canonical cell cycles have been described: endoreplication, in which cells undergo G and S phases but do not initiate M phase; and endomitosis, where cells execute all four cell-cycle phases but do not initiate or complete cytokinesis during M phase.

Endoreplication and endomitosis both result in the generation of polyploid cells that are widespread in animals and plants and play important functions in the growth of tissues and organs, the generation of nutrients and in fine-tuning gene expression^7^.

Previous research on polyploid cells in insects, plants and mice has unveiled several aspects of the mechanisms by which cells switch from canonical cell cycles to endoreplication. The developmental cues and signalling pathways that drive endoreplication are generally tissue- specific, but they converge on the downregulation of mitotic CDK (M-CDK) activity, which is required for entry into M phase^8^. For example, in the mouse trophectoderm lineage, accumulation of the CDK inhibitor proteins p57^KIP^^2^ and p21^CIP^^1^ makes cells bypass M phase and transition to G1 phase, driving endoreplication and differentiation into trophoblast giant cells^9^. In *Drosophila melanogaster* ovaries, endoreplicating follicle cells block M-phase entry by inhibiting the expression of String, a Cdc25-type phosphatase that is required for M-CDK activation^10^. Furthermore, artificial inhibition of M-CDK activity can induce endoreplication in many different cell types and organisms^11–13^. Thus, inhibition of M-CDK activity seems to be a general mechanism by which cells can transition from canonical to endoreplication cycles.

In contrast to the regulation of endoreplication cycles, much less is known about how cells transition to endomitosis cell cycles. As endomitosis cycles are defined by the presence of an M phase, cells must have sufficient M-CDK activity to enter M phase. Interestingly, cells can perform endomitosis in different manners: some cells, such as polyploid mammalian megakaryocytes, enter M phase but exit at an early stage, largely before chromosome segregation, leading to mononucleated polyploid cells^14^. In contrast, mammalian hepatocytes proceed further into M phase, segregating chromosomes but failing to undergo cytokinesis, thus becoming binucleated^15–18^. It is unclear what mechanisms allow some endomitotic cells to progress further in M phase than others, and how cells can uncouple nuclear division from cytokinesis, two processes that are normally tightly coupled, as they are both initiated by the dephosphorylation of CDK1 substrates and destruction of mitotic proteins by the anaphase- promoting complex (APC)^19^. One possible mechanism of uncoupling M phase and cytokinesis could be a reduced activity or expression of M phase proteins specifically involved in cytokinesis. Indeed, altered expression of cytokinesis regulators has been reported for some mammalian cell types that undergo endomitosis^14,17,20,21^. For example, murine megakaryocytes downregulate the RhoA guanine exchange factors GEF-H1 and ECT2, resulting in impaired cytokinesis^14^. Strikingly, overexpression of GEF-H1 and ECT2 in diploid megakaryocytes is sufficient to prevent both endomitosis and the emergence of polyploid megakaryocytes. Thus, an altered expression of cytokinesis regulators could underly the transition from canonical to endomitosis cell cycles. It remains to be determined, however, whether repression of cytokinesis regulators is a general mechanism to initiate endomitosis cycles, and if so, how different endomitotic cell types achieve this repression.

There is some evidence that the inhibition of cytokinesis during endomitosis could be regulated through changes in transcription. For example, the downregulation of GEF-H1 during megakaryocyte polyploidization involves the transcriptional cofactor Mkl1^14,22^. Furthermore, in rodents, the atypical E2F transcription factors E2F7 and E2F8 are required to transition to endomitosis cell cycles in hepatocytes by repressing cytokinesis genes that are normally activated by E2F1 during canonical cell cycles^18,23^. However, both Mkl1 and the atypical E2Fs regulate a wide range of processes other than the cell cycle by controlling the expression of many genes, including genes involved in cell differentiation^23,24^. Thus, it is currently unclear how specific changes in cell-cycle gene expression that ultimately either promote endomitosis or endoreplication are achieved during development^25,26^. Studying how cells transition between different cell-cycle types remains a significant challenge, as these cell-cycle transitions are not easily recapitulated *in vitro*, nor are they easily tractable in complex model organisms such as mice, zebrafish, and fruit flies, that consist of hundreds of thousands of somatic cells. An ideal model system would offer the ability to predict exactly when cells undergo endomitosis and allow their study at single-cell resolution *in vivo.* The intestinal lineage of the nematode *Caenorhabditis elegans* meets these criteria due to its simplicity and invariant cell-cycle pattern: intestinal cells perform canonical cell cycles during embryogenesis and then perform one endomitosis cell cycle at the end of the first larval stage (L1) thereby becoming binucleated, followed by an endoreplication cycle at the end of each larval stage (L1 to L4) which increases the final cellular ploidy to 64C (**Figure 1A**)^27–29^.

**Figure 1.**
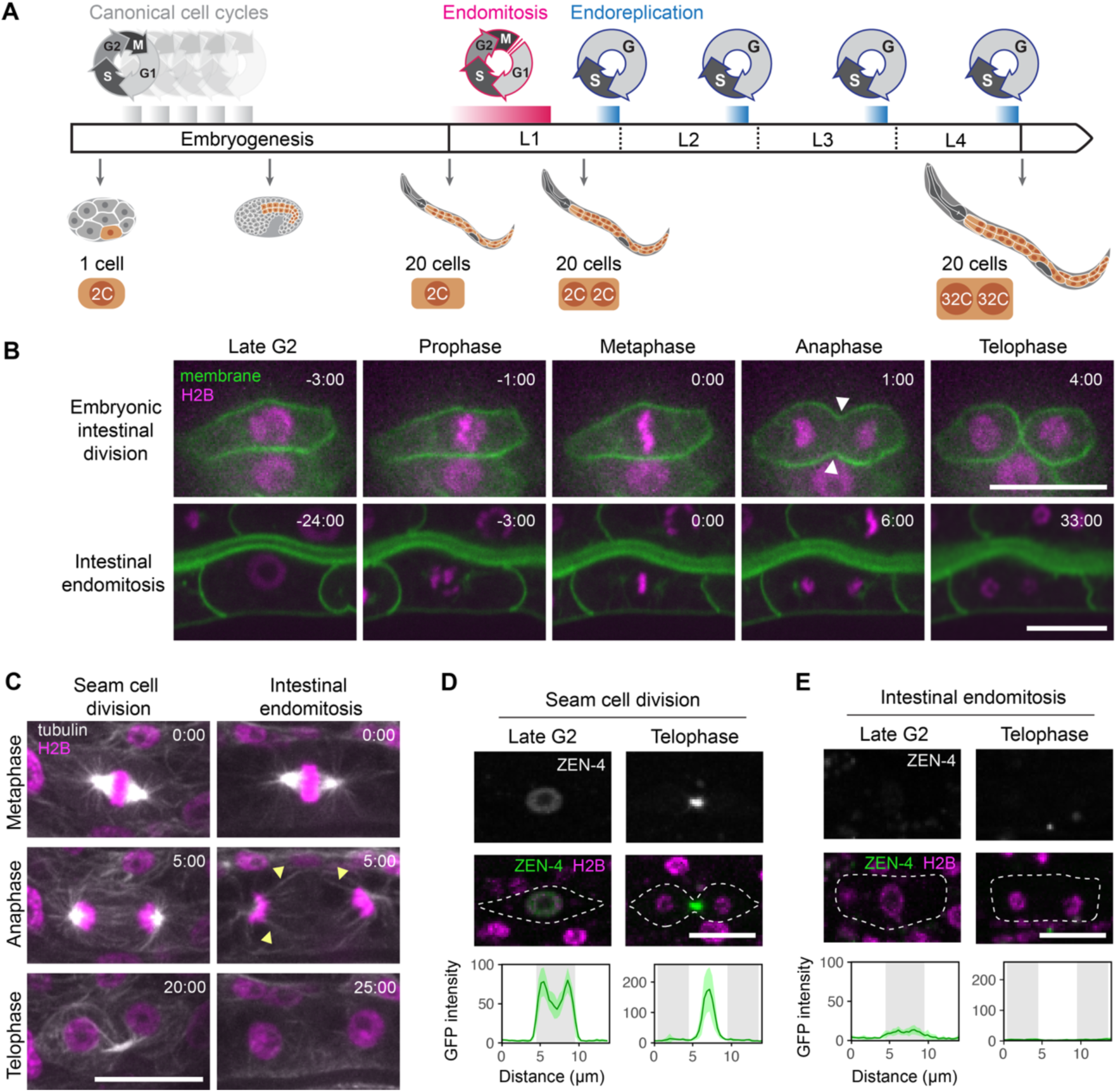
Intestinal cells do not form a central spindle or initiate cytokinesis during endomitosis. **(A)** Schematic depicting intestinal development and cell-cycle transitions across embryogenesis and larval development. In early embryogenesis, intestinal cells undergo multiple canonical cell cycles, after which a functional intestine is formed. During the first larval stage (L1), the 14 most posterior intestinal cells (from intestinal rings 3-9) undergo one endomitosis cycle that gives rise to binucleated cells. Shortly thereafter, intestinal cells transition to endoreplication, which takes place at the end of each larval stage (L1-L4). **(B)** Stills from live imaging of intestinal cells during canonical cycles in embryogenesis (top) and endomitosis M phase in L1 larvae (bottom) in a strain expressing GFP-PH (green) and H2B-mCherry (magenta) to mark intestinal cell membranes and DNA, respectively. Stills represent different stages of M phase, where 0:00 (min:sec) is the frame before anaphase is first observed (defined as “metaphase”). White arrows mark the formation of cleavage furrow (n ≥ 15 for all conditions, N = 3 experiments). **(C)** Representative images of canonical M phase in seam cells (left) and endomitosis M phase in intestinal cells (right) from live imaging of a strain expressing H2B-mCherry (magenta) and tubulin-GFP (gray) in all somatic cells. Stills depict different stages of M phase, starting from metaphase (0:00, min:sec). Yellow arrowheads point to microtubules present in the cell midzone during anaphase of intestinal endomitosis. (n ≥ 18 for all conditions, N = 2 experiments). **(D-E)** Representative images of seam and intestinal cells in late G2 phase **(D)** or in telophase **(E)** of M phase during canonical or endomitosis M phase, respectively. Animals express endogenously tagged GFP::ZEN-4 (green) and ubiquitously expressed H2B-mCherry (magenta), and intensity profile plots on the bottom depict GFP::ZEN-4 fluorescence intensities centered around the DNA (gray background) during G2 phase and in telophase. Images are maximum projections of 3-5 slices. Mean intensity is shown by the dark green line, standard deviation (SD) is represented in light green (n ≥ 21 for all condi(ons, N = 2 experiments). Scale bars depict 10 μm in all images.

Here, we study how endomitosis is regulated in the *C. elegans* intestine and show that intestinal cells do not initiate membrane furrowing and lack a central spindle during endomitosis M phase. We find that many M-phase genes are more lowly expressed during intestinal endomitosis. Among these, genes encoding central spindle factors are most profoundly affected and, in contrast to other cell-cycle genes, these cytokinesis genes also show strongly reduced protein levels. We find that intestinal cells lose the capacity to undergo cytokinesis after late embryogenesis, suggesting that the cell-cycle transition to endomitosis is a developmentally programmed switch. Furthermore, we find that the evolutionarily conserved DREAM complex contributes to the repression of cytokinesis genes and the timely transition to endomitosis during development. Taken together, our findings indicate that transcriptional downregulation of cytokinesis genes drives the cell-cycle transition to endomitosis in the *C. elegan*s intestine, providing novel insights into the mechanisms by which cells modify their cell cycles to become polyploid during development.

## RESULTS

### Cytokinesis is not initiated during intestinal endomitosis

To understand how cytokinesis is inhibited in intestinal cells during endomitosis M phase, we first compared M phases of canonical versus endomitosis cycles during intestinal development. We used a reporter strain expressing GFP-PH and H2B-mCherry markers in the intestinal lineage to mark cell membranes and nuclei, respectively, and performed live imaging of intestinal cells undergoing either canonical M phases during embryogenesis or endomitosis M phases during the first larval stage. As expected, embryonic intestinal cells undergo all steps of M phase, starting with DNA condensation and nuclear envelope breakdown (NEB), followed by chromosome segregation, cleavage furrow ingression and completion of cytokinesis (**Figure 1B**, **Movie S1**). In contrast, intestinal cells undergoing endomitosis M phase exhibit normal DNA condensation, NEB and chromosome segregation, but do not initiate membrane ingression during M-phase exit (**Figure 1B**, **Movie S2**). Thus, cleavage furrowing is not initiated during intestinal endomitosis, suggesting that cytokinesis is inhibited at an early stage.

Cleavage furrow formation and ingression are mainly directed by the mitotic spindle during anaphase, when different populations of microtubules signal to the cell cortex to induce furrowing^30,31^. Because the central spindle is essential for initiating cytokinetic furrowing in many animal cells, we wondered whether the early inhibition of cytokinesis during endomitosis could be explained by the absence of the central spindle. Therefore, we performed live imaging of a strain expressing GFP-tubulin and H2B-mCherry to visualize the mitotic spindle during canonical and endomitosis M phases. In both cases, a bipolar mitotic spindle could be observed during prometaphase and metaphase, and astral microtubules were seen to emanate from the centrosomes towards the cell midzone and cortex during anaphase (**Figure 1C**). However, as opposed to canonical M-phase, microtubules did not form a detectable central spindle structure during endomitosis M phase, likely explaining the failure to initiate cytokinesis during endomitosis.

We then wondered whether the lack of central spindle assembly could be explained by the absence of the proteins that are normally required for its formation. Many of the proteins needed for central spindle assembly, such as Aurora B and Cdk1, are also essential for earlier events of M phase such as bipolar spindle formation and chromosome segregation. As we did not observe any defects during early M phase, it is unlikely that these proteins are absent during endomitosis. One protein complex that is specifically required for central spindle assembly and cytokinesis, but not for the earlier events of M phase, is the centralspindlin complex consisting of ZEN-4 (KIF23, also known as MKLP1) and CYK-4 (RACGAP1)^31^. To investigate the centralspindlin complex, we endogenously tagged the ZEN-4^MKLP^^1^ subunit with GFP in a strain expressing H2B- mCherry in all somatic cells, allowing us to visualize ZEN-4^MKLP^^1^ expression during canonical and endomitosis M phases. In seam cells undergoing canonical cell cycles, ZEN-4^MKLP^^1^ localizes to the nucleus during late interphase, then relocates to the mitotic spindle and cell membrane during early M phase and to the midzone region during anaphase where it concentrates at the midbody during late telophase (**Figure 1D**). Analysis of intestinal cells revealed that ZEN-4^MKLP^^1^ levels are remarkably lower during late interphase of endomitosis cycles and ZEN-4^MKLP^^1^ levels remain low throughout endomitosis M phase (**Figure 1E**). Taken together, these experiments reveal that intestinal cells lack a central spindle and have strongly reduced expression of the centralspindlin complex component ZEN-4^MKLP^^1^, likely explaining the absence of cleavage furrow formation and cytokinesis during endomitosis.

### Cytokinesis genes are transcriptionally downregulated during intestinal endomitosis

Reduced ZEN-4^MKLP^^1^ levels during endomitosis could be a consequence of either post- transcriptional regulation, for example through decreased mRNA or protein stability, or by altered *zen-4* transcription. To understand the apparent absence of ZEN-4^MKLP^^1^ during endomitosis we first investigated *zen-4* mRNA levels using single molecule fluorescence in situ hybridization (smFISH)^32^. Cell-cycle regulated genes are typically expressed in two major waves of gene expression that peak during the G1/S and G2/M cell-cycle transitions, and are hereafter referred to as G1/S and G2/M genes, respectively^33^. Genes involved in cytokinesis are classified as G2/M genes, hence we examined the *zen-4* mRNA levels during M phase^34^. We compared *zen- 4* levels during intestinal endomitosis with *zen-4* levels in seam cells undergoing canonical cell cycles (**Figure 2A**). In endomitosis M phase, *zen-4* mRNA levels are approximately four-fold lower than in canonical M phase, suggesting that reduced ZEN-4 protein levels during endomitosis M phase are due to decreased mRNA expression (**Figure 2B**). To understand whether other genes involved in cytokinesis are also downregulated during intestinal endomitosis, we analyzed the mRNA levels by smFISH of the other centralspindlin complex component *cyk-4*, the microtubule crosslinker *spd-1* (*PRC1*), and components of the contractile ring *nmy-2* (*myosin II*), *ect-2* (*ECT2*) and *let-502* (*ROCK*)^31^. Similarly to *zen-4*, mRNA levels of *cyk-4* and *spd-1* are also strongly downregulated during endomitosis M phase. Furthermore, the expression of the contractile ring regulators *nmy-2*, *ect-2* and *let-502* is also significantly reduced (**Figure 2C**). Collectively, these results indicate a general downregulation of multiple cytokinesis genes during endomitosis, and a particularly strong reduction in the expression of the central spindle regulators *zen-4*, *cyk-4* and *spd-1*.

**Figure 2.**
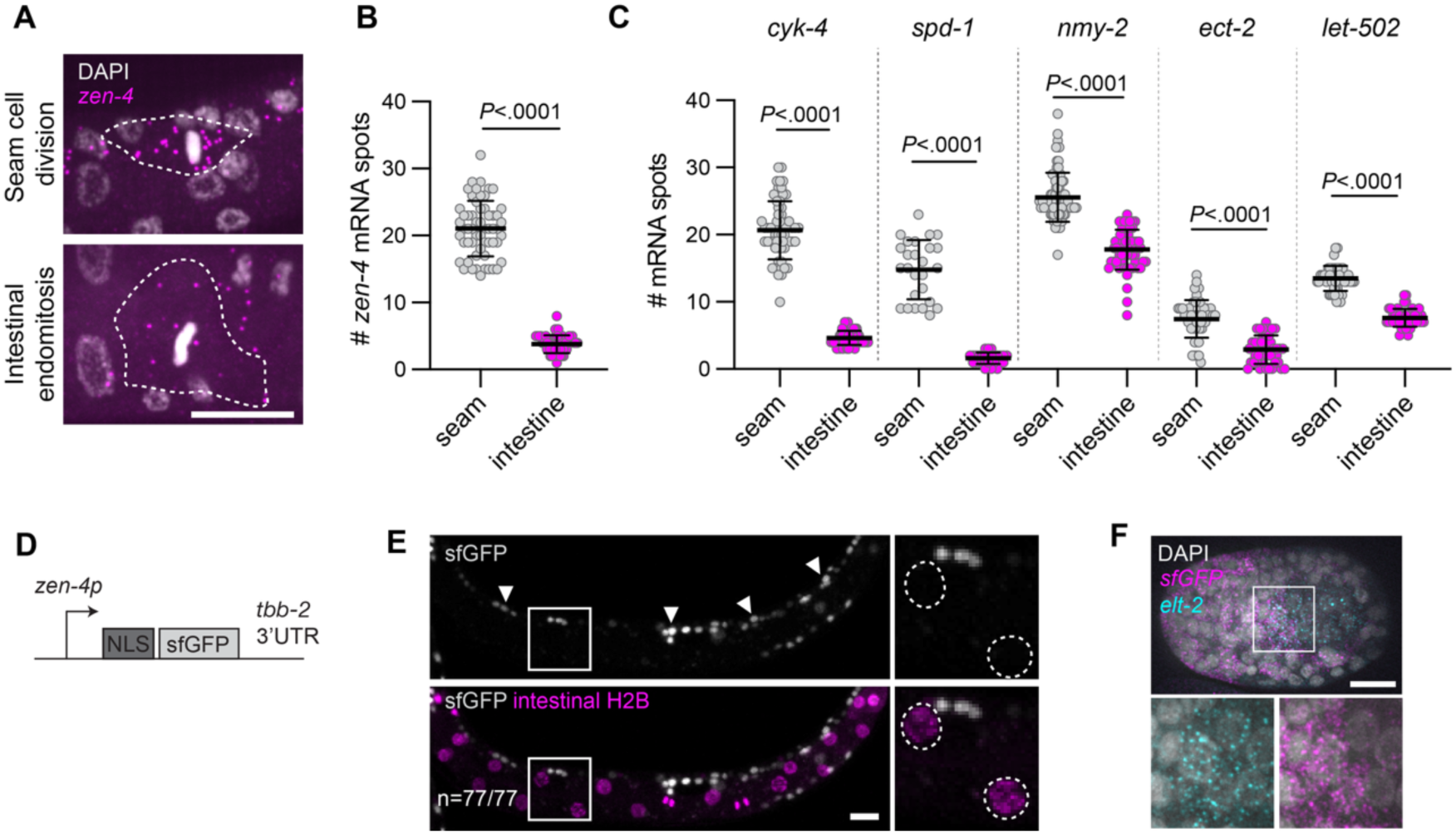
Intestinal endomitosis is characterized by the transcriptional downregulation of multiple essential cytokinesis genes. **(A)** Representative images of smFISH showing DAPI staining (grey) and *zen-4* mRNA spots (magenta) during metaphase of a canonical seam cell division (top) and intestinal endomitosis M phase (bottom). Dashed lines represent the cell membranes marked by GFP-PH (not shown). Images are maximum projections of 12 slices. **(B)** Quantification of number of *zen-4* mRNAs per cell during M phase of seam cell divisions and intestinal endomitoses (n ≥ 52 cells, , N ≥ 2). The black lines and error bars indicate the mean ± SD. *P* value was calculated by Mann-Whitney test. **(C)** Quantifications of number of *cyk-4*, *spd-1*, *nmy-2*, *ect-2* and *let-502* mRNAs per cell during M phase of seam cell divisions and intestinal endomitoses (n ≥ 26 cells for all conditions, N ≥ 2). The black lines and error bars represent the mean ± SD. *P* values were calculated by Mann-Whitney test. **(D)** Schematic representation of the *zen-4p::^NLS^sfGFP* transgene (*matIs192*) in which a 648 bp fragment of the *zen-4* promoter drives the expression of NLS-sfGFP. **(E)** Representative images of a late L1 larva carrying the *zen-4p::^NLS^sfGFP* transgene (grey) and expressing H2B-mCherry (magenta) in intestinal cells. Arrowheads indicate non-intestinal cell nuclei. Close-up images show two GFP-negative intestinal nuclei (dashed circles). Images are a maximum projection of 9 slices (n = 77 animals, N = 3 experiments). **(F)** Representative image of smFISH for *sfGFP* (magenta) and *elt-2* (cyan) mRNAs and DAPI staining (grey) in an embryo carrying the *zen-4p::^NLS^sfGFP* transgene. Bottom images are close-ups of the same region of the embryo, with intestinal cells containing *sfGFP* mRNA molecules. Close-up images show a region with intestinal cells (n = 26, N = 2 experiments). Scale bars depict 10 µm in all images.

Low mRNA levels of cytokinesis genes could be a consequence of either reduced transcription, or by altered mRNA stability. To test whether cytokinesis gene expression levels in intestinal cells are controlled at the transcriptional or post-transcriptional level, we generated a transcriptional reporter for *zen-4* expression. A multicopy array containing a 648 bp promoter fragment upstream of the *zen-4* start codon driving the expression of a nuclear GFP (*zen- 4p::^NLS^sfGFP*) was integrated in a strain expressing nuclear mCherry in intestinal cells (**Figure 2D**). Analysis of L1-stage larvae revealed multiple GFP-positive nuclei that, based on their localization, correspond to seam cells, cells of the P lineage, mesoblast progenitors and the germline, all of which undergo canonical cell cycles and divide during larval development (**Figure 2E**)^27^. However, we did not find any intestinal cells with detectable nuclear GFP, indicating that the activity of the *zen-4* promoter is remarkably low during intestinal endomitosis (n= 77/77 animals, **Figure 2E**). Thus, reduced *zen-4* mRNA levels during intestinal endomitosis are likely due to a lack of transcription.

Early embryonic cell cycles are largely driven by maternally provided mRNAs^35^. This raises the possibility that intestinal cells have sufficient maternally provided *zen-4* to perform canonical cell cycles during embryogenesis, but never activate *zen-4* transcription, leading to the absence of ZEN-4^MKLP^^1^ protein during endomitosis in the first larval stage. smFISH analysis of *zen- 4* transcripts in embryos revealed high levels of *zen-4* expression in one- to eight-cell stage embryos, likely reflecting maternally provided transcripts (**Figure S1A**). We then examined the expression of the *zen-4p::^NLS^sfGFP* reporter in embryos using fluorescent probes against *sfGFP*. Due to germline silencing of the multicopy *zen-4p::^NLS^sfGFP* reporter, *sfGFP* mRNAs are not maternally loaded into embryos and could only be detected from the four-cell stage onwards, when embryonic transcription starts (**Figure S1B)**. Strikingly, all somatic cells of embryos between the eight-cell stage and the 32-cell stage (when intestinal cells are performing canonical cell cycles) contained many cytoplasmic *sfGFP* mRNAs molecules as well as bright nuclear transcriptional foci (**Figure S1C**), suggesting that intestinal progenitor cells are expressing *zen-4*. Furthermore, we could observe *sfGFP* mRNA molecules in cells that also express the intestine- specific gene *elt-2* (**Figure 2F**), indicating that intestinal cells are able to activate *zen-4* expression during canonical cell cycles. As expected, *zen-4p::^NLS^sfGFP* expression in late embryos is absent in most cells, as they have entered quiescence (**Figure S1D**). Taken together, our results suggest that intestinal cells lose the capacity to transcribe cytokinesis genes after the completion of their canonical cell cycles during embryogenesis, resulting in the absence of cytokinetic ingression during endomitosis M phase.

### mRNA expression of many G2/M genes is reduced during endomitosis

We next employed genome-wide mRNA sequencing to investigate whether additional genes are differentially expressed during endomitosis. As mentioned earlier, cytokinesis gene expression is usually restricted to dividing cells, and is triggered together with many other G2/M genes during a major gene-expression wave that begins during S phase and peaks at the G2/M transition^34^. Thus, to understand whether other G2/M genes are downregulated during endomitosis, we isolated cells in S/G2/M phase of endomitosis or canonical cell cycles by fluorescence-associated cell sorting (FACS) from a strain expressing intestine-specific mCherry (*elt-2p::mCherry-PH-P2A-H2B-mCherry*). Synchronized L1 larvae were grown for six hours to allow intestinal cells as well as cells of other lineages to enter the cell cycle (**Figure 3A**). We sorted 500 mCherry-positive or mCherry-negative cells with a 4N DNA content based on Hoechst dye intensity and performed bulk RNA sequencing (**Figure 3B and Figure S2**). The non-intestinal cell population is therefore a mixture of different cell types that normally divide during the L1 stage. Transcriptional profiles of the different samples confirmed that intestinal and non-intestinal cell populations clustered separately (**Figure S3A**). Differential gene expression analysis showed 1254 and 1138 genes significantly enriched with intestinal and non-intestinal cells, respectively (Log2FC > 2, padj < 0.05, **Figure 3C**). As expected, tissue-enrichment analysis of genes significantly associated with the intestinal cell population reveals the intestine as the first most enriched tissue (**Figure S3B**), whereas genes enriched with the non-intestinal cell population show anatomical terms related to the germline or Q-cell lineage (**Figure S3C**). Consistent with our smFISH experiments, the cytokinesis genes *zen-4*, *cyk-4*, *spd-1, ect-2* and *nmy-2* are expressed significantly higher in non-intestinal cells, supporting the notion that they are expressed at lower levels during endomitosis (**Figure S4A** and **Figure 3C**).

**Figure. 3.**
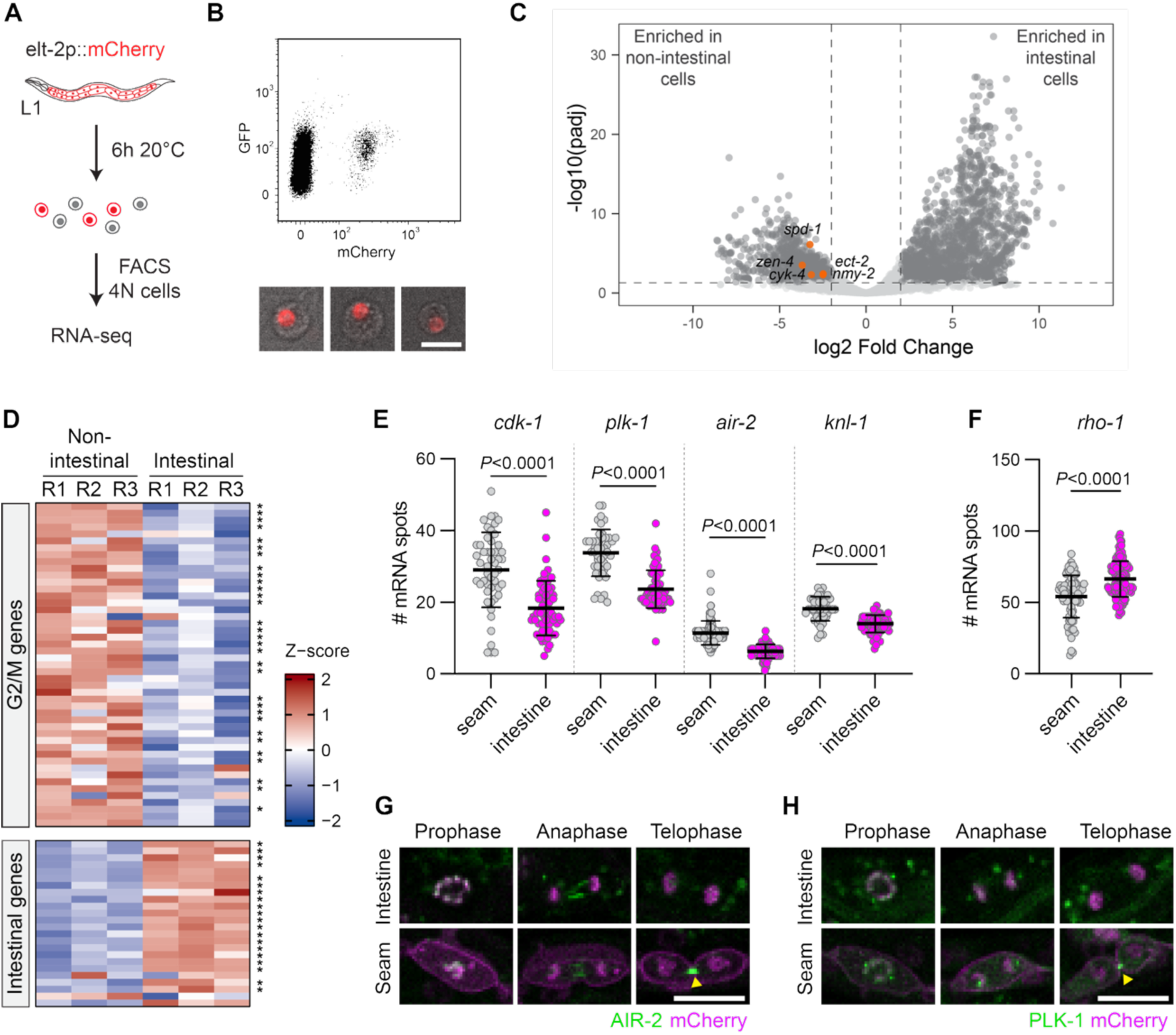
mRNA expression levels of G2/M cell-cycle genes is lower during endomitosis compared to canonical cell cycles. **(A)** Schematic representation of intestinal cell purification and isolation for RNA sequencing experiments. L1 larvae of a strain expressing H2B-mCherry and mCherry-PH in the intestine were fed for six hours prior worm dissociation and cell sorting of 4N cells by FACS. **(B)** Dot plot showing the GFP and mCherry levels of single cells derived from the worm dissociation (top) and example images of isolated mCherry-positive intestinal cells (bottom). **(C)** Volcano plot showing differential gene expression between intestinal (mCherry-positive) and non-intestinal (mCherry-negative) cells. Dashed lines represent the padj < 0.05 and log2FC > -2|2 cutoffs. Cytokinesis genes *zen-4*, *cyk-4*, *spd- 1*, *ect-2* and *nmy-2* (highlighted in orange) are enriched in the non-intestinal cells. **(D)** Heatmap showing the Z-score and the log2FC of a subset of G2/M genes (top) or intestinal-specific genes (below) Asterisks indicate statistical significance (padj < 0.05 and log2FC < -2 for G2/M genes; padj < 0.05 and log2FC > 2 for intestinal-specific genes). **(E)** smFISH quantifications of number of *cdk-1*, *plk- 1*, *air-2* or *knl-1* mRNAs per cell in intestinal endomitosis or seam cell M phases (n ≥ 43 for all conditions, N ≥ 2 experiments). The black lines and error bars represent the mean ± SD. *P* values were calculated by Mann-Whitney test. **(F)** smFISH quantification of *rho-1* mRNA spots in intestinal endomitosis and seam cell cycles at M phase (n ≥ 83, N = 3 experiments). The black lines and error bars correspond to the mean ± SD. *P* value was calculated by Mann-Whitney test. **(G-H)** Representative images of endogenous PLK-1 **(G)** and AIR-2 **(H)** protein localizations during different stages of M phase during intestinal endomitosis (top) or seam cell division (bottom). Intestinal nuclei are marked by H2B-mCherry (magenta), and seam cell membranes and nuclei are marked by mCherry-PH (magenta) and H2B-mCherry (magenta), respectively. Yellow arrows mark the localization of PLK-1 or AIR-2 at the cytokinetic bridge. Scale bar depicts 10 µm in all images.

We manually curated a list of *C. elegans* G2/M genes to find whether they are differentially expressed during endomitosis. Interestingly, 84.8% of G2/M genes are expressed significantly higher in the non-intestinal cell population, suggesting that the expression of a large portion of cell-cycle regulated genes is reduced during endomitosis (**Figure 3D**). Similarly, many G1/S genes are also expressed significantly higher in non-intestinal cells (**Figure S4B**). Among the many G2/M genes that show downregulation in intestinal cells, we found key mitotic factors that are required for M-phase entry and progression (**Figure S4C**). smFISH analyses of *cdk-1* (*CDK1*), *plk-1* (*PLK1*), *air-2* (*AURKB*) and *knl-1* (*KNL1*) expression in either endomitotic or canonical M phases confirmed the mRNA sequencing results, pointing to a reduced expression of many M phase genes during endomitosis (**Figure 3E**). While *cdk-1*, *plk-1*, *air-2* and *knl-1* showed reduced expression in endomitosis, the fold reduction in expression levels between canonical and endomitosis M phases was smaller than for the cytokinesis genes *zen-4, cyk-4* and *spd-1* (**Figure 3E and Figure 2B,C**). In addition, the expression of *rho-1*(*RhoA*), a small GTPase that plays an important role in cytokinesis but is not expressed in a cell-cycle dependent manner in mammalian cells^36^, was not downregulated in intestinal cells (**Figure 3F**). These results suggest that most G2/M regulated genes are expressed at lower levels during endomitosis, with central spindle genes being downregulated most strongly.

### Mitotic protein abundance and localization during endomitosis M phase are not altered

Most G2/M genes, including *cdk-1*, *plk-1* and *air-2*, are crucial for cell division and have early functions during M phase. Thus, it is striking that these genes are downregulated during endomitosis, given that intestinal cells enter M phase and perform normal nuclear divisions. We therefore hypothesized that mitotic factors must be present at sufficient levels at the protein level, despite being downregulated at the transcriptional level. To check the protein levels of mitotic factors during endomitosis, we crossed strains with endogenously tagged PLK-1::GFP and AIR-2::GFP with strains containing intestine or seam cell-specific membrane and nuclear markers. Similar to what has been previously described in mammalian cells^37^, PLK-1^PLK1^ localizes at the centrosomes and DNA throughout prophase until anaphase, and to the cytokinetic bridge during anaphase and telophase during canonical divisions in seam cells. We did not notice apparent changes in PLK-1^PLK1^ abundance or localization during endomitosis M phase, except that PLK-1^PLK1^ fails to localize to the midzone during telophase (**Figure 3G**). AIR-2^AURKB^ is also expressed at similar levels in both intestinal and seam cells, where it localizes on the DNA during prophase. Consistent with its role at the central spindle^38^, AIR-2^AURKB^ then re-localizes to the midzone during anaphase in seam cells, and finally concentrates at the cytokinetic bridge during telophase. In contrast, AIR-2^AURKB^ does not concentrate at the midzone during anaphase and telophase of endomitosis in intestinal cells (**Figure 3H**). Nevertheless, we observed GFP signal in filamentous structures at the midzone during anaphase, indicating that AIR-2^AURKB^ can interact with astral microtubules at the interpolar region despite the absence of a central spindle (**Figure 1B**). Taken together, our results indicate that although G2/M genes are transcriptionally downregulated during endomitosis, the essential mitotic proteins AIR-2^AURKB^ and PLK-1^PLK1^ are expressed at apparently normal levels and localize to their expected subcellular locations (except for the central spindle and cytokinetic bridge, which are absent during endomitosis M phase, see above).

### Intestinal cells are unable to express high levels of cytokinesis genes after embryogenesis

Our results suggest that during development, intestinal cells downregulate the expression of cytokinesis regulators, resulting in a lack of cytokinetic furrowing during endomitosis M phase. We wondered whether intestinal cells lose the ability to undergo canonical cell cycles during a specific moment in development. To investigate when intestinal cells lose the capacity to perform cytokinesis, we generated a strain carrying an intestine-specific transgene expressing *cye-1* (cyclin E) and a gain-of-function allele of *cdk-2* (CDK2), which have previously been shown to force ectopic divisions in mammalian cells as well as in *C. elegans*^39–41^. By examining newly-hatched L1 larvae, we found that intestinal overexpression of CYE-1 and CDK-2^AF^ during embryogenesis resulted in many additional intestinal cell divisions, but also gave rise to binucleated cells, suggesting endomitosis can already occur during embryogenesis (**Figure 4A** and **B**). To test whether intestinal cells are able to perform cytokinesis after embryogenesis, we generated a strain with a heat-shock inducible promoter driving the expression of CYE-1 and gain-of-function alleles of CDK-2 and CDC-25.1 (Cdc25), which are sufficient to drive additional M phases in *C. elegans* larvae^41^. We hypothesized that overexpressing these cell-cycle regulators would increase G2/M gene expression, thereby potentially boosting cytokinesis gene expression and driving cytokinesis. However, we found that induction of CYE-1, CDK-2^AF^, and CDC-25.1^GF^ in L2 larvae gives rise to additional nuclear divisions in the intestine (**Figure 4C-E**), but does not give rise to additional intestinal cells (N=42/42, **Figure 4F**). This was not due to an inability of the CYE- 1, CDK-2^AF^, and CDC-25.1^GF^ proteins to induce cytokinesis, because similar overexpression experiments in starved larvae resulted in ectopic division of seam cells (N=17/20, **Figure 4G**). Together, these results indicate that intestinal cells lose the capacity to undergo canonical cell cycles at some point during embryogenesis after which cytokinesis cannot be induced even when forcing ectopic cell cycles.

**Figure 4.**
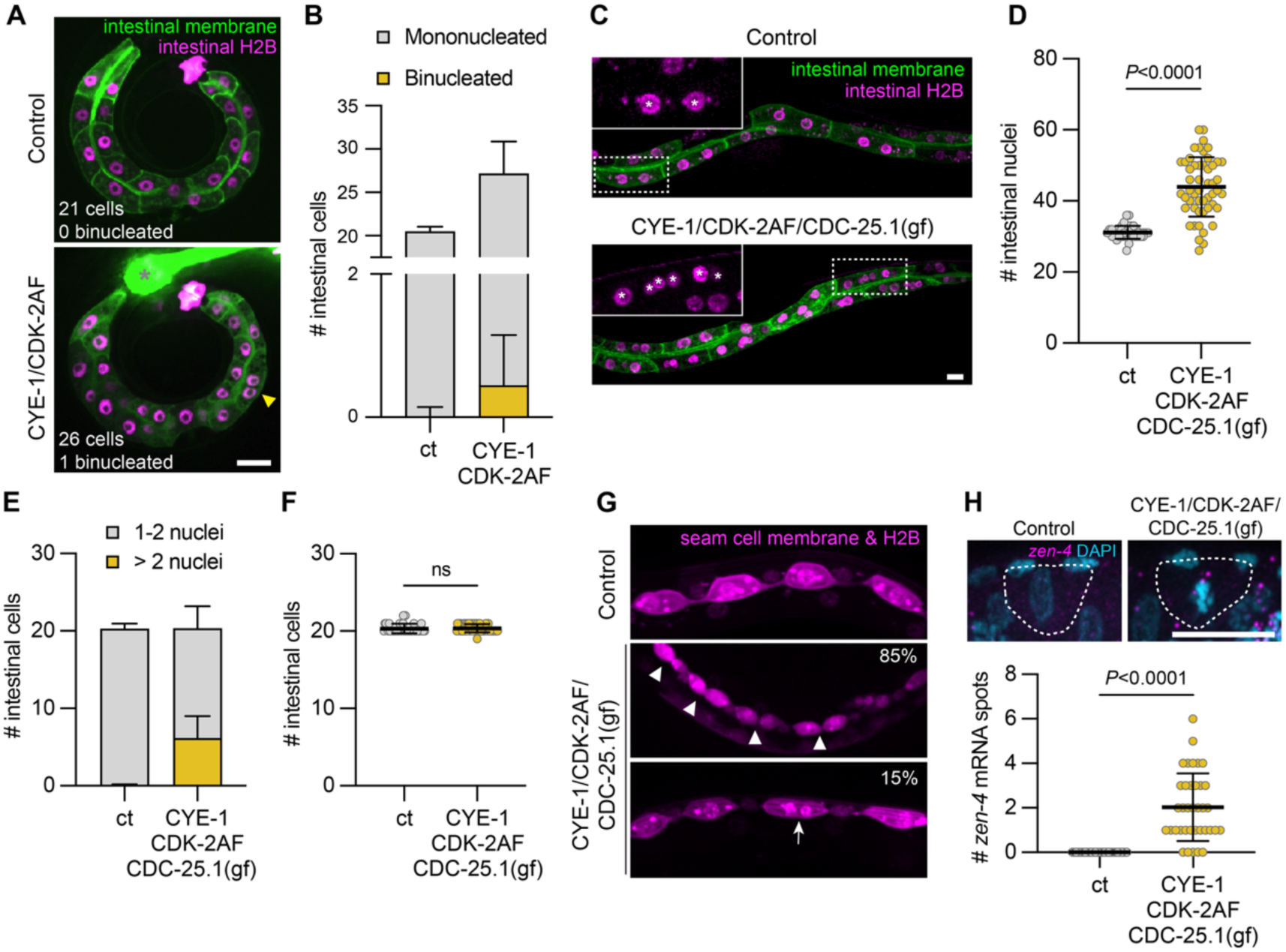
The cell-cycle transition to endomitosis is a robust switch induced at the end of embryogenesis. **(A)** Representative images of early L1 larvae with GFP-PH (green) and H2B-mCherry (magenta) to mark intestinal membranes and nuclei from a control strain (top) and a strain overexpressing CYE- 1/CDK-2AF in intestinal cells (bottom). Yellow arrowhead highlights a binucleated cell. The gray asterisk marks the myo-2*p-GFP* co-injection marker that is present in the CYE-1/CDK-2AF overexpression strain. **(B)** Average number of mononucleated and binucleated cells per worm in early L1 larvae from a control (ct) strain and a strain with intestinal-specific expression of CYE-1 and CDK-2AF (n ≥ 57 for all conditions, N = 2). Error bars indicate the SD. **(C)** Representative images of L3 larvae after heat-shock in control (ct) or a strain with heat-shock inducible expression of CYE- 1, CDK-2AF and CDC-25.1(gf). Close-ups show an intestinal cell with nuclear mCherry marker (H2B- mCherry) in magenta, white asterisks depict each nucleus of the cell. **(D-E)** Average number of intestinal nuclei per worm **(D)** or per cell **(E)** after heat shock at the L2 stage in control (ct) and the heat-shock inducible CYE-1/CDK-2AF/CDC-25.1(gf) overexpression strain (n ≥ 42 for all conditions, N = 2 experiments). Horizontal lines and error bars represent mean ± SD. The *P* value was calculated by Mann-Whitney test. **(F)** Average number of intestinal cells per worm in heat-shocked animals of the control strain (ct) or the CYE-1/CDK-2AF/CDC-25.1(gf) heat-shock inducible overexpression strain. (n ≥ 42 for all conditions, N = 2 experiments). Horizontal lines and error bars represent mean ± SD. The *P* value was calculated by Mann-Whitney test. **(G)** Representative images of early L1 stage animals expressing seam cell membrane and nuclear markers (magenta) in heat-shocked control (ct) and CYE-1/CDK-2AF/CDC-25.1(gf) overexpression strains. The two images in CYE- 1/CDK-2AF/CDC-25.1(gf) illustrate different cell-cycle phenotypes. White arrowheads point to seam cells undergoing cytokinesis, and arrow shows a binucleated seam cell (n ≥ 20 animals per condition, N = 2 experiments). **(H)** Representative images of *zen-4* smFISH in control (ct) or after heat-shock induced overexpression of CYE-1/CDK-2AF/CDC-25.1(gf). Quantification of number of *zen-4* mRNAs per cell is shown below (n ≥ 37 per condition, N = 2 experiments). Horizontal lines and error bars represent mean ± SD. The *P* value was calculated by Mann-Whitney test. Scale bar depicts 10 µm in all images.

The inability of larval intestinal cells to perform cytokinesis, even when they are forced into additional M phases, could be explained by their inability to express cytokinesis genes during the G2/M wave of cell-cycle gene expression. Alternatively, it is possible that ectopic cell cycles do increase cytokinesis gene expression, but that there are additional mechanisms that supress cytokinetic furrowing during endomitosis. To test whether cytokinesis genes are expressed after induction of ectopic M phases, we analysed the expression of *zen-4* by smFISH. For these experiments, we induced ectopic M phases in early L1 stage animals by heat-shock activated expression of CYE-1, CDK-2^AF^, and CDC-25.1^GF^ in starved L1 stage animals, and then fed animals for six hours at 20 degrees before fixing them for smFISH analyses. At this stage intestinal cells are normally in S/G2 phase of endomitosis, and indeed all control animals (which were heat shocked but did not contain the heat shock-inducible CYE-1, CDK-2^AF^, and CDC-25.1^GF^ transgenes) had mononucleated intestinal cells and did not show any signs of M-phase entry (**Figure 4G**). In contrast, animals with overexpressed CYE-1, CDK-2^AF^, and CDC-25.1^GF^ contained many intestinal cells in M phase (**Figure 4G**). Strikingly, also in these ectopic M phases, *zen-4* transcripts remain low, similar to what we observe during the normal endomitosis M phase that occurs at the late L1 stage (**Figure 4H** and **2B**). Thus, the induction of ectopic M phases is not sufficient to substantially activate the expression of *zen-4*, suggesting that cytokinesis genes become stably repressed during larval intestinal development.

Repression of cytokinesis genes could be mediated by the formation of heterochromatin at the regulatory elements of these genes. To investigate whether chromatin landscape of G2/M genes is altered in cells undergoing endomitosis, we mapped various histone modifications associated with active (H3K4me3) or repressive chromatin (H3K27me3 and H3K9me3) using a chromatin immunocleavage sequencing (ChIC-seq) protocol adapted for low cell numbers^42^. We isolated and fixed cells from one-hour old L1 larvae from a strain expressing intestinal-specific mCherry markers, followed by Hoechst staining and FACS to purify G1-phase populations of intestinal and non-intestinal cells based on the mCherry signal and DNA content. As expected, we find that intestinal-specific genes such as *act-5* and *asp-5* are enriched for the active H3K4me3 mark in intestinal cells, and for the repressive H3K27me3 mark in non-intestinal cells, confirming the validity of our approach (**Figure 5A**). However, we did not find notable differences in the tested histone modifications on the *zen-4* and *spd-1* loci between intestinal and non- intestinal cells (**Figure 5A**). In addition, G2/M genes exhibit low levels of H3K27me3 and H3K9me3 in both intestinal and non-intestinal cells (**Figure 5B**). Mechanistically, these results suggest that the reduced expression of G2/M genes during endomitosis is not caused by the deposition of H3K27me3 or H3K9me3 repressive marks in their genomic loci.

**Figure 5.**
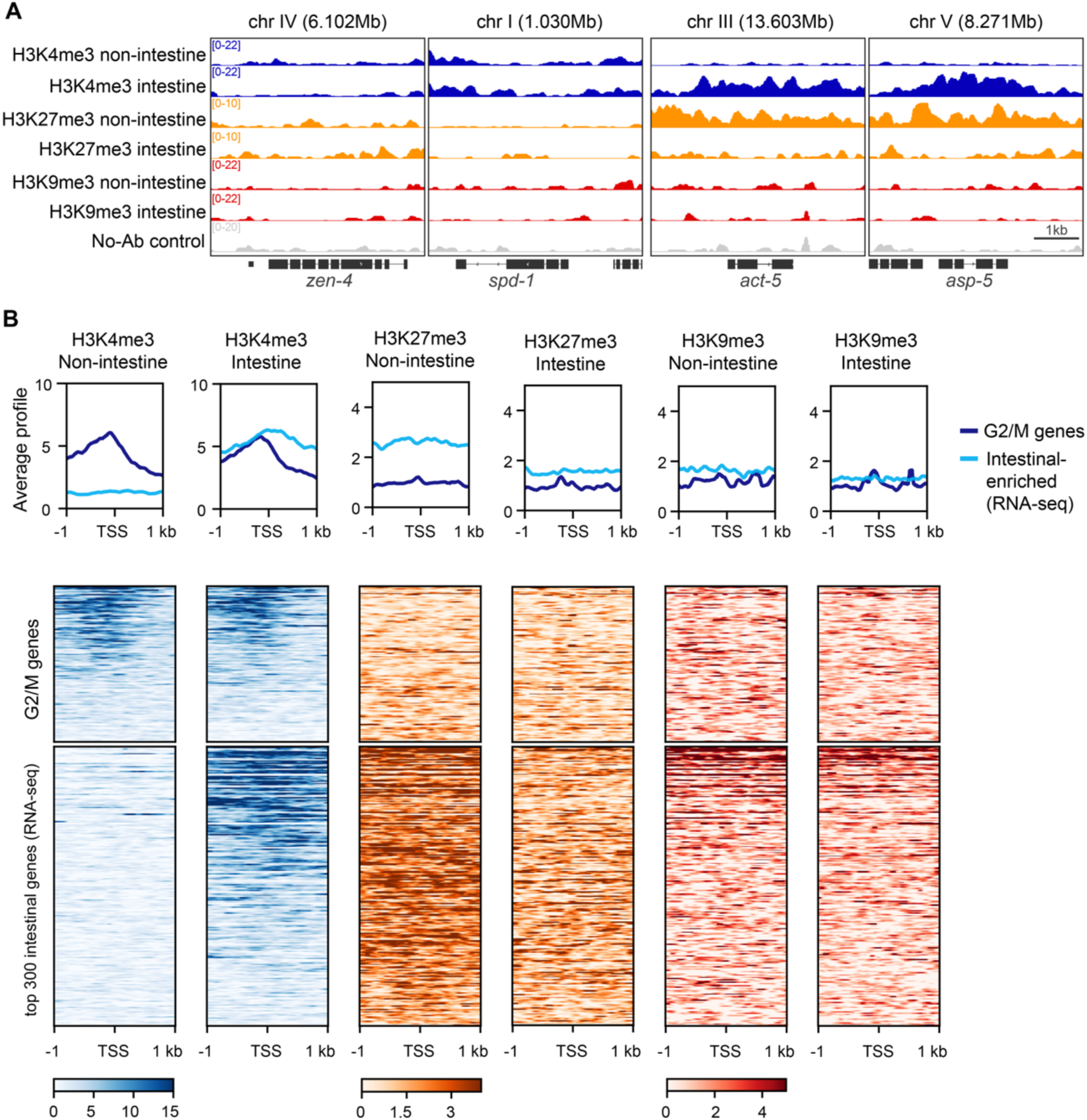
G2/M gene loci are not enriched for H3K27me3 or H3K9me3 repressive marks in endomitosis. **(A)** Genomic regions containing the *spd-1*, *zen-4*, *act-5* and *asp-5* loci showing the H3K4me3 (blue), H3K27me3 (orange) and H3K9me3 (red) ChIC-seq signal. **(B)** Average plots and heatmaps of H3K4me3, H3K27me3 and H3K9me3 ChIC-seq coverage in non-intestinal and intestinal cells over a set of G2/M genes and the 300 genes most enriched intestinal cells (fold change) from the RNA-seq dataset. In the heatmaps, each row represents one gene locus from 1 kb upstream until 1 kb downstream of the transcription start site (TSS).

### The DREAM complex is involved in the repression of cytokinesis genes during endomitosis

To elucidate the mechanism by which cytokinesis genes are repressed during endomitosis, we investigated whether conserved regulators of G2/M gene expression are important for cytokinesis gene repression. In animals, the timely expression of G2/M genes is controlled by the conserved DREAM complex (<u>D</u>P, <u>R</u>B-related, <u>E</u>2F <u>a</u>nd <u>M</u>uvB), which represses the expression of G2/M genes early in G1 phase and during quiescence^43^. In mammals, rising CDK activity has been shown to disrupt the DREAM complex, releasing the MuvB complex that can then interact with the transcription factors B-MYB and FOXM1 to activate the expression of G2/M genes^33,44^. In *C. elegans,* DREAM consists of the DPL-1/EFL-1 heterodimer, the pocket protein LIN-35 and the MuvB complex (LIN-9, LIN-37, LIN-52, LIN-53 and LIN-54), however, there are no clear orthologs of B-MYB or FOXM1 in *C. elegans* making it unclear how G2/M gene expression is activated^45,46^. To test whether DREAM is important to repress cytokinesis genes during intestinal endomitosis, we investigated whether depletion of the DREAM complex members DPL-1, EFL-1, LIN-9 or LIN-54 by RNAi could activate the *zen-4* transcriptional reporter in the intestine. Strikingly, 50% of *efl-1* RNAi animals and 85% of *dpl-1* RNAi animals showed detectable nuclear GFP expression in their intestinal cells, compared to 0% of animals treated with the control RNAi *lon-1* (**Figure 6A** and **B**). RNAi knockdown of *lin-54* and *lin-9* also resulted in intestinal *zen-4p::^NLS^sfGFP* expression, albeit in a smaller fraction of animals (**Figure 6A** and **B**). Notably, although RNAi depletion of *dpl-1*, *efl-1*, *lin-9* or *lin-54* led to the expression of the *zen- 4p::^NLS^sfGFP* reporter, nuclear GFP levels in intestinal cells were generally lower than in cells undergoing canonical cell cycles, suggesting that inhibition of DREAM is not sufficient to completely activate *zen-4* expression during endomitosis. To verify a role of DREAM in the repression of cytokinesis genes during endomitosis, we quantified the expression of endogenous z*en-4* during endomitosis M phase by smFISH in *efl-1*, *dpl-1* and *lin-54* mutants. Because null mutants of these genes are not viable, we used a temperature sensitive allele of *efl-1(se1)*, and *lin-54(n3423)* or *dpl-1*(gk685) homozygous mutants that arise from heterozygous *dpl- 1(gk685)/mIn1* and *lin-54(n3423)/nT1* animals. We found a significant increase in the expression of *zen-4* during endomitosis upon depletion of *efl-1* or *dpl-1*, and a modest but significant increase in *lin-54* mutants (**Figure 6C**). Despite the increased mRNA levels, *zen-4* levels remained below the levels observed in seam cells undergoing a canonical M phase (**Figure 2B**). Of note, the *dpl-1 and lin-54* mutants likely received maternal *dpl-1 and lin-54* mRNAs from their heterozygous mothers, which could result in residual DPL-1 or LIN-54 during early larval development of homozygous mutants, and could possibly explain the limited extent of *zen-4* re- expression. Nonetheless, we tested whether the observed increase in zen*-4* expression in DREAM mutants is sufficient to drive cytokinesis during endomitosis M phase by analysing the number of intestinal cells at the L2 larval stage in *dpl-1* and *lin-54* mutants, just after endomitosis has completed. If depletion of DREAM would lead to a conversion of endomitosis cell cycles into canonical divisions, we would detect more than the wild-type 20 cells in the intestine (depending on how many cells would be able to divide, the number of cells could vary between 20 and 40). We counted the number of intestinal cells in L3 stage *dpl-1(gk685)* and *lin-54(n3423)* mutants that had been treated with either control (*lon-1*), *dpl-1* or *lin-54* RNAi, to exclude possible redundancy or contribution from maternal products. We found no differences in the number of intestinal cells between control and *dpl-1* and *lin-54* mutants, indicating cytokinesis was not induced during endomitosis (**Fig 6D**, *dpl-1(gk685)* and *lin-54(n3423)* mutants grown on the control RNAi). Furthermore, RNAi of *dpl-1* did not alter the number of intestinal cells in *dpl- 1(gk685)* nor *lin-54(n3423)* mutants. However, *lin-54* RNAi gave rise to additional intestinal cells in both *dpl-1(gk685)* and *lin-54(n3423)* mutants (**Figure 6D**). The presence of additional intestinal cells upon depletion of *lin-54* could either be explained by the induction of cytokinesis during endomitosis M phase, or alternatively, by additional rounds of canonical cell cycles during embryogenesis. To distinguish between these possibilities, we quantified the number of intestinal cells in newly hatched L1 larvae. We found that both *dpl-1(gk685)* and *lin-54(n3423)* mutants treated with *lin-54* RNAi were born with more than 20 intestinal cells, showing that DREAM is required to restrict the number of canonical cell cycles during embryogenesis (**Figure 6E**). Thus, despite a role of the DREAM complex in the repression of cytokinesis genes during endomitosis, there are likely additional mechanisms that further repress cytokinesis genes to ensure that cells do not divide.

**Figure 6.**
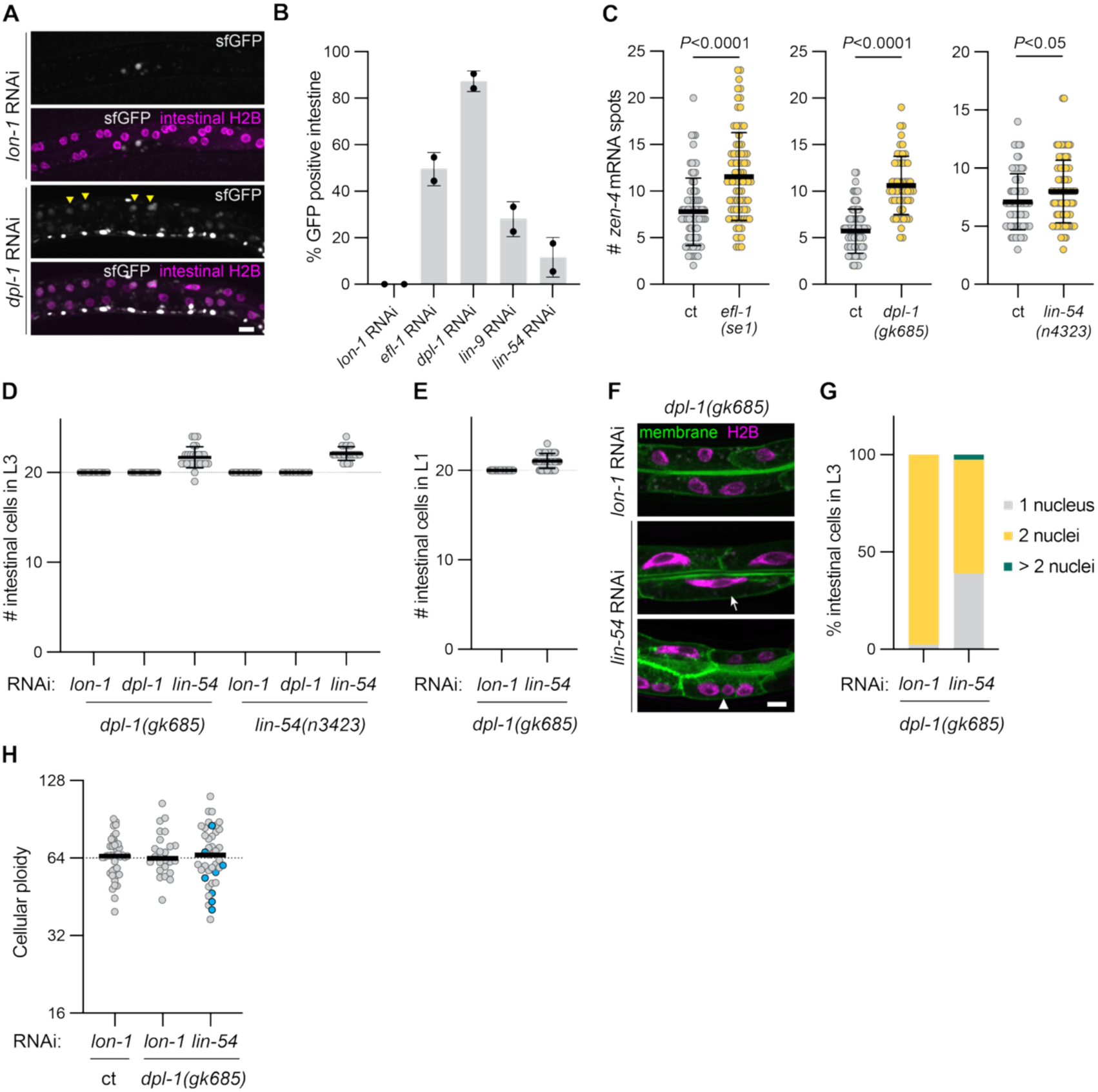
The DREAM complex is involved in the repression of *zen-4* during endomitosis and promotes the transition to endomitosis cycles during embryogenesis **(A)** Representative images of animals expressing *zen-4p::^NLS^sfGFP* (grey) and intestinal H2B- mCherry after RNAi depletion of *lon-1* (control) or *dpl-1*. Yellow arrowheads mark GFP-positive intestinal nuclei after depletion of *dpl-1*. **(B)** Percentage of worms with at least one GFP-positive intestinal cell in control (*lon-1*) or *efl-1*, *dpl-1*, *lin-9* or *lin-54* RNAi-treated animals carrying the *zen- 4p::^NLS^sfGFP* transgene. Each point represents the average percentage of animals with a GFP- positive intestine per experiment in each condition and error bars indicate the standard deviation (n ≥ 8 animals per condition, N = 2 experiments). **(C)** smFISH quantifications of number of *zen-4* mRNAs per cell during intestinal endomitosis M phase in control or *efl-1(se1)* mutants grown at restrictive-temperature (25°C), *dpl-1* heterozygous control (*ct)* or homozygous (*dpl-1(gk685)*) mutants, and *lin-54* heterozygous controls (*ct*) or homozygous (*lin-54(n3423)*) mutants (n ≥ 54 in all conditions, N = 3 experiments. Horizontal lines and error bars represent mean ± SD. The *P* value was calculated by Mann-Whitney test. **(D)** Average number of intestinal cells per worm at the L3 stage in *dpl-1(gk685)* and *lin-54(n3423)* homozygous mutants treated with *lon-1*, *dpl-1* or *lin-54* RNAi. Horizontal lines and error bars represent mean ± SD. The *P* value was calculated by Mann- Whitney test. **(E)** Average number of intestinal cells per worm in newly hatched L1 larvae from homozygous *dpl-1(gk685)* mutants upon control or *lin-54* RNAi treatment. Horizontal lines and error bars represent mean ± SD. The *P* value was calculated by Mann-Whitney test. **(F)** Representative images of intestinal cells in *dpl-1(gk685)* homozygous mutants treated with *lon-1* or *lin-54* RNAi. The two images in *dpl-1(gk685)* + *lin-54* RNAi show the different phenotypes observed: mononucleated cells with elongated nuclei (white arrowhead) and intestinal cells with more than two nuclei (white arrow). **(G)** Percentage of intestinal cells with one, two or more than two nuclei in homozygous *dpl- 1(gk685)* mutants treated with *lon-1* or *lin-54* RNAi (n ≥ 41, N = 2 experiments). **(H)** Quantification of intestinal cell ploidy of adult stage *dpl-1* heterozygous control animals treated with the control lon-1 RNAi and *dpl-1(gk685)* homozygous mutants treated with *lon-1* or *lin-54* RNAi. Gray data points represent binucleated cells, and blue data points are mononucleated cells. Only intestinal cells of the Int3 ring were analyzed (n ≥ 24 for all conditions, N = 2 experiments). Horizontal lines indicate the median. Dashed line represents the wild type Int3 cellular ploidy (64C). All scale bars depict 10 μm.

### Depletion of DREAM components converts endoreplication to endomitosis cycles

In addition to an increased intestinal cell number, we noticed that *dpl-1(gk685)* animals treated with *lin-54* RNAi contained many mononucleated cells with elongated nuclei, reminiscent of cells that had failed mitosis, as well as cells with more than two nuclei, which are never present in wild-type animals (**Figure 6F** and **G**). These nuclear alterations after depletion of DREAM could be a result of cells performing additional (and potentially aberrant) cell cycles during larval development, similar to the additional canonical divisions that we observed after DREAM depletion during embryogenesis. Alternatively, it is possible that the DREAM complex is more directly involved in the regulation of non-canonical cell cycles, and its depletion is converting endoreplication cycles, which normally occur at the end of every larval stage (**Figure 1A**), into endomitosis, leading to cells with more than two nuclei. To determine whether loss of LIN-54 and DPL-1 results in additional cell cycles or alterations in the type of cell cycle, we measured the ploidy of intestinal cells in adult animals. Heterozygous *dpl-1(gk685)*/*mIn1* animals and their progeny were grown on RNAi feeding plates for five days, after which adults were fixed for quantitative DNA staining to measure cellular ploidies. As expected, RNAi knockdown of *lin- 54* resulted in 100% sterile progeny. Ploidy measurements of intestinal cells in *dpl-1(gk685)* animals treated with *lin-54* RNAi showed no significant increases in cellular ploidy, showing that depletion of DREAM does not give rise to additional non-canonical cell cycles (**Figure 6H**). Thus, intestinal cells with more than two nuclei are likely arising by the conversion of endoreplication cycles into endomitosis, rather than by additional cell cycles. Altogether, these data points to a role of the DREAM complex in regulating the cell-cycle switch from endomitosis to endoreplication, likely by enabling a progressive repression of G2/M gene expression during non-canonical cell cycles (**Figure S5)**.

## DISCUSSION

Alterations to the canonical cell cycle are widespread in multicellular organisms and ensure the correct formation of tissues and organs during development. A growing number of studies has brought new insights in many cell-cycle alterations that occur during development, in particular endomitosis and endoreplication, two non-canonical cell cycles that give rise to somatic polyploidy^47^. Although the mechanisms that cells use to initiate and perform endoreplication have been largely elucidated, much less is known on how endomitosis is regulated and whether there is a common mechanism that allows endomitotic cells to inhibit cytokinesis. In this study, we investigated *C. elegans* intestines to understand how cells initiate endomitosis and are able to uncouple nuclear division from cellular division. Our results uncover a global repression of cell-cycle genes during endomitosis M phase, suggesting that cells undergo a transcriptional reprogramming of their cell-cycle genes when they transition to non-canonical cell cycles. We find that cytokinesis genes become robustly repressed during endomitosis, and cannot become re-expressed even after forcing additional M phases during larval development. Our results suggest that the conserved DREAM repressor complex functions redundantly to repress cytokinesis genes during endomitosis, ensuring the correct transition from canonical to non-canonical cycles during development.

Through live imaging, we observed that intestinal embryonic cell cycles undergo all stages of canonical M phase. In contrast, M phase of intestinal endomitosis is characterized by a nuclear division in the absence of cleavage furrowing. This complete absence of cytokinetic furrowing has also been described for murine hepatocytes undergoing endomitosis, but is not observed in human hepatocytes or murine cardiomyocytes, where cells undergo cleavage furrow ingression before aborting cytokinesis^18,48,49^. Our analyses also revealed that intestinal cells do not form a central spindle, which could explain the absence of cytokinetic furrowing in endomitosis M phase. However, not all endomitotic cells that do not undergo cytokinetic furrowing also lack a central spindle^14,50^. Therefore, it seems like there are multiple means by which cells abort cytokinesis during endomitosis, but it remains unclear if different organisms have developed independent strategies to inhibit specific cytokinetic events or whether common mechanisms can explain the transition to endomitosis across species.

Despite the apparent diversity in how cells undergo endomitosis in different organisms, there are also similarities. Endomitosis often occurs during clear developmental transitions: hepatocytes become binucleated after weaning, rat cardiomyocytes after embryogenesis and mammary glands during pregnancy^15,51,52^. Similarly, we find that *C. elegans* cells undergo a clear transition from canonical cell cycles to endomitosis during late embryogenesis. This cell-cycle switch seems very robust, considering that forcing cells into a cell cycle after late embryogenesis will only give rise to additional nuclei but not additional cells. In addition, our study revealed another potential commonality between different cell types that undergo endomitosis, namely a transcriptional reprogramming of cell-cycle gene expression. Similar to the cytokinesis gene expression changes that have been reported in mammalian hepatocytes and megakaryocytes, we find that intestinal cells reprogram their cell-cycle gene expression during endomitosis.

We demonstrate that the mRNA levels of cytokinesis genes are notably decreased during M phase of intestinal endomitosis, compared to canonical M phases in seam cells. Of note, we observed that the expression of genes encoding for central spindle components (*zen-4*, *cyk-4* and *spd-1*) shows a considerably larger fold reduction than that of cytokinesis genes involved in contractile ring assembly (*nmy-2*, *ect-2*, *let-502*). What the underlying reason is for these differences in expression levels is currently unclear. In addition to low mRNA levels of cytokinesis regulators, we also find that protein levels of the centralspindlin complex member ZEN-4^MKLP1^ are substantially reduced during endomitosis M phase. Consistent with the absence of ZEN-4^MKLP1^, intestinal cells in endomitosis M phase fail to form a central spindle, which is essential to concentrate key cytokinesis regulators such as ECT2 at the division plane, thus potentially explaining the absence of cytokinesis initiation in endomitosis^53^. Interestingly, we do observe interpolar microtubules that are enriched with AIR-2^AURKB^, suggesting that the inability to form a central spindle is most likely due to the absence of the centralspindlin complex or SPD-1^PRC1^, rather than the inability to recruit AIR-2^AURKB^ or a lower stability of interpolar microtubules. Since multiple cytokinesis factors (e.g. ZEN-4^MKLP1^, CYK-4^RACGAP1^ and SPD-1^PRC1^) show reduced expression, loss of cytokinesis can likely not be attributed to the altered expression of a single gene, and therefore rescuing cytokinesis through ectopic expression of one or more factors is likely challenging.

Despite the fact that intestinal cells in endomitosis perform nuclear divisions, our RNA sequencing analysis revealed that many G2/M genes are expressed significantly lower during endomitosis compared to canonical cell cycles. These findings could be influenced by differences inherent to comparing transcriptomes of different cell lineages; however, we believe this is not the case because the expression of cell-cycle genes during specific phases of the cell cycle is usually invariable between different cell types^33^. Indeed, we were able to validate by smFISH that several G2/M genes were expressed at lower levels during endomitosis M phase compared to seam cell M phases. This reduced expression of G2/M phase genes is intriguing, considering that many of them encode factors that are essential for M-phase entry and chromosome segregation, both of which occur normally in endomitosis M phase. We can envision two scenarios by which endomitotic cells can undergo M phase with reduced G2/M gene expression: one possibility is that mitotic protein levels are reduced but are still sufficient to allow cells to perform nuclear division; alternatively, lowered mRNA levels of mitotic factors do not result in lower abundance of their encoded proteins. Our analyses of the M-phase factors PLK-1^PLK1^ and AIR-2^AURKB^ revealed that they are found abundantly at the protein level, in stark contrast to ZEN-4 ^MKLP1^, whose levels are hardly detectable. We also find that mRNA levels of *zen-4* are more reduced than mRNA levels of *plk-1* and *air-2*, which could imply that there is a specific threshold in transcript abundance that allows sufficient protein translation, and that endomitotic cells can produce enough levels of M-phase proteins necessary for nuclear division, but are unable to synthesize cytokinesis factors like ZEN-4^MKLP1^.

The DREAM complex plays an important role in the timely expression of G2/M genes by repressing their expression during earlier cell cycle phases and in quiescence^33^. Our analyses revealed an increase in *zen-4* mRNA levels in DREAM complex component mutants (*efl-1*, *dpl-1* and *lin-54* mutants), indicating that the DREAM complex contributes to the repression of cytokinesis genes during intestinal endomitosis. These findings are consistent with existing whole-organism microarray and RNA-sequencing data sets from *C. elegans* which, despite lacking cell-type resolution, show that cytokinesis genes are globally upregulated in DREAM mutants^54–56^. Decreased expression of DREAM targets in endomitosis could indicate that the mechanisms that normally relieve G2/M gene repression by DREAM in M phase are absent in intestinal cells. In mammalian cell cycles, expression of G2/M genes begins upon the disassembly of DREAM and the recruitment of transcriptional activators B-Myb and FoxM1 to the MuvB complex^33^. However, it is unknown how the activation of G2/M gene expression occurs in *C. elegans* as no orthologs of B-Myb and FoxM1 have been described, despite that the B-Myb interacting domains of the MuvB complex are evolutionarily conserved in *C. elegans*^57,58^. Thus, identifying the alternative mechanisms that enable G2/M gene expression in *C. elegans* will help us understand how G2/M gene expression is dampened during intestinal endomitosis.

Despite the fact that *zen-4* mRNA levels in *efl-1, dpl-1* and *lin-54* mutants are increased in endomitosis M phase, they are still on average two-fold lower than *zen-4* mRNA levels in canonical cell cycles in seam cells. These results suggest that there are additional mechanisms that inhibit cytokinesis gene expression during endomitosis M phase. Indeed, DREAM inactivation does not induce cytokinesis during intestinal endomitosis, which would be expected if the endomitosis cycle is converted into a canonical cell cycle. Therefore, it is likely that other factors in addition to DREAM, cooperate in the repression of cytokinesis genes during endomitosis. Our results suggest that the downregulation of cytokinesis as well as other G2/M genes is not mediated by the enrichment of H3K27me3 or H3K9me3, histone modifications commonly associated with heterochromatin. Nonetheless, it remains possible that specific histone modifications or histone variants are deposited at G2/M genes loci during endomitosis, such as the histone variant H2A.Z that was shown to be enriched on DREAM targets^55^.

An involvement of the DREAM complex in the regulation of non-canonical cell cycles has also been reported in plants and in *Drosophila*^59–61^. In *Arabidopsis* leaves, the switch from canonical to endoreplication cycles is delayed upon RNAi knockdown of *E2FA*, which is similar to the phenotype we observe upon *lin-54* knock down in the *C. elegans* intestine^59^. In *Drosophila*, loss of *dDP* in the fat body, a tissue that normally undergoes endoreplication, results in increased ploidy and the appearance of multinucleated cells^60^, suggesting that cells are converting endoreplication cycles into endomitosis cycles, a phenotype we also observe upon DREAM knockdown in *C. elegans*. It is remarkable that such diverse species as plants, nematodes and insects may employ similar mechanisms to switch from canonical to non-canonical cycles during development, and it will be interesting to investigate whether DREAM complex is also involved in the regulation of endomitosis in mammals.

## METHODS

### Worm culture and strains

*C. elegans* were cultured on nematode growth medium (NGM) plates at 20°C according to standard protocols, unless indicated otherwise. Worms were fed with OP50 *E. coli* bacteria, except for RNA sequencing and RNAi experiments (see below).

Strains used in this study are listed in **Table S1.** The intestine-specific membrane (*matIs46)* and H2B-mCherry (*ccyIs27)* transgenes were made using the miniMos based single- copy insertion method with a plasmid containing *ges-1p::myrGFP* or *elt-2p::mCherry-PH-P2A- H2B-mCherry* sequences, respectively^62^. The *zen-4p::^NLS^sfGFP* reporter (*matIs192)* was made by gamma-irradiating a strain carrying an extrachromosomal array with a plasmid containing the 648 bp promoter upstream of the *zen-4* start codon fused with the *egl-13* nuclear localisation sequence (NLS), *sfGFP* and the *tbb-2* 3’UTR, as well as the *lin-48p::TdTomato* and *sur-5p::mCherry* co-injection markers. The transgene *ccyIs12* was made by integrating a single copy of the plasmid *dpy-30p::H2B::mCherry::P2A::sfGFP* using the mos1-mediated single copy insertion (mosSCI) method as previously described^63^. Endogenously tagged *GFP::zen-4* was generated using CRISPR/Cas9-mediated homologous recombination using a self-excising selection cassette approach as previously described^64^.

### RNA interference

Feeding RNAi experiments were performed as previously described using HT115 clones from the Vidal RNAi library, carrying the corresponding target gene in a L4440 vector ^65^. To induce the production of dsRNA, bacteria were grown overnight at 37°C in LB medium supplemented with Ampicillin and Tetracyclin, then diluted in the morning 1:1 with fresh LB supplemented with 2 mM IPTG (final concentration 1 mM), incubated one hour at 37°C, then concentrated 10-fold and seeded onto NGM plates supplemented with IPTG and Ampicillin. L4 larvae from the corresponding genotypes were transferred the RNAi NGM plates and were grown for four days at 20°C until the next generation were gravid adults. Animals were then washed off the plates and newly hatched L1 larvae were collected after an hour and transferred onto new RNAi NGM plates until the desired stage.

For RNAi depletions of *efl-1*, *dpl-1*, *lin-9*, *lin-54* and *lon-1* in the strain expressing *zen- 4p::^NLS^sfGFP*, double-stranded RNA (dsRNA) was injected into the gonads of young-adult stage animals. Injected animals were transferred to new plates after 24 and 48 hours, and L2 larvae derived from eggs that were laid after 48 hours post-injection were analysed by microscopy.

### Microscopy and image analysis

Static imaging of ZEN-4::GFP, PLK-1::GFP and AIR-2::GFP strains was performed on a Leica SPE confocal microscope. For imaging, animals were mounted onto microscopy slides with 2% agarose pads and immobilized with 1-2 mg/ml of levamisole. For imaging of ZEN-4::GFP, PLK- 1::GFP and AIR-2::GFP and the membrane and/or nuclear mCherry markers, L1 larvae were synchronized by wash-off, and grown for 18-20 (late L1, intestinal endomitosis) or 22-24 hours (early L2, seam cell division) at 15°C. Animals were imaged using a 40x objective and a slice thickness of 0.5 µm. Imaging of DREAM mutants and RNAi experiments was performed on a on a Nikon Ti2 microscope equipped with a CSU-X1 spinning disk scanner unit and a CMOS digital camera (C13440; Hamamatsu Photonics, Hamamatsu-city, Japan) using a 40x objective and 2x binning. For static imaging of animals after heat shock, worms were mounted on a 2% agarose pad containing 30 mM of NaN3, and images were acquired with a Leica SPE confocal microscope and a 40x oil objective.

Time-lapse live imaging was performed on a Nikon Ti2 microscope equipped with a CSU- X1 spinning disk scanner unit and a CMOS digital camera (C13440; Hamamatsu Photonics, Hamamatsu-city, Japan), using a 100x objective and 2x binning. For imaging the embryonic intestinal cell cycles, embryos were collected by splaying gravid adults and were mounted on slides containing a 2% agarose pad. For imaging the intestinal endomitosis M phase, L1 larvae were synchronized by bleaching gravid adult animals and leaving embryos to hatch overnight in M9 buffer, and were grown for 20 to 22 hours at 15°C. For imaging of TBA-2::GFP, L1 larvae were synchronized by wash-off, and grown for 20 to 24 hours at 15°C prior mounting on a 2% agarose pad containing levamisole.

For quantitative DNA staining, young adult hermaphrodites were fixed with 4% PFA and stored in 70% ethanol at 4°C overnight. DAPI staining was performed using 5 μg/ml DAPI in 1 ml wash solution (10% formamide, 2× SSC) for 60 minutes at 37°C, after which animals were mounted on a slide using ProLong gold antifade mountant (Invitrogen #P36934). Imaging was performed on either a Nikon Ti2 microscope equipped with a CSU-X1 spinning disk or a Leica SPE confocal microscope using a 40x objective and 0.5 µm Z-slices.

All image analysis was performed using FIJI^66^. Quantifications of ZEN-4 levels were performed using the plot profile tool: a line was drawn on top of the chosen cell, and the pixel intensity from a single Z-slice (where the DNA was the brightest) along the distance of the line was measured. For ploidy measurements, z-projections were made of the Z-slices containing the nuclei of interest and integrated densities of the nuclear regions were measured. For binucleated cells, the integrated densities of the 2 nuclei were summed. For each animal, at least three control (2N) pharynx or body wall muscle nuclei were also measured, and their average nuclear intensities were used to calculate the ploidy of the measured intestinal cells.

### Heat-shock experiments

All heat-shock experiments were carried out at 33°C for either 30 or 60 minutes. For measuring the effects of CYE-1, CDK-2^AF^, and CDC-25.1^GF^ overexpression in intestinal cells, L1 larvae were synchronized by wash-off and grown on NGM plates for 20 hours at 15°C. Plates were then placed in a 33°C water bath for 30 minutes, then transferred to 15°C and imaged 25 hours later. To overexpress CYE-1, CDK-2^AF^, and CDC-25.1^GF^ in seam cells, heat shocks were performed on starved L1 larvae in M9 buffer in 1.5 ml tubes for one hour. Heat-shocked animals were grown on NGM plates for 3 hours at 15°C. For the smFISH experiments in intestinal cells upon overexpression of CYE-1, CDK-2^AF^, and CDC-25.1^GF^, starved L1 larvae were heat shocked in M9 buffer in 1.5 ml tubes for one hour and grown on NGM plates for 5.5 hours at 20°C prior to fixation.

### Single molecule fluorescence in situ hybridization (smFISH)

smFISH was performed as previously described ^32^. To facilitate quantification, we made use of transgenic animals expressing membrane markers (GFP-PH) in either intestinal cells (alleles *matIs53* and *matIs46*) or seam cells (allele *heIs166*). Animals were fixed at the desired developmental stage using 4% PFA and stored in 70% ethanol for up to two months. Hybridization was carried out for 16-18 hours at 37°C. Oligonucleotide probes complementary to the mRNA of the corresponding gene were designed using the Stellaris probe designer (https://www.biosearchtech.com/) and chemically labelled with the fluorescent dyes atto565 or atto633 as previously described^67^. Samples were washed and stained with DAPI prior to mounting. Z-stacks of 0.2 µm thickness were generated using a spinning disk confocal microscope equipped with C11440-22C camera, using a 100x oil objective and 2x binning. mRNA spots were manually counted in Fiji using the cell counter tool. Only fluorescent spots present in more than 2-3 Z-slice s and within the cell boundaries were counted.

### Worm dissociation and intestinal cell isolations by FACS

For intestinal cell isolations, liquid cultures were grown of 200,000-400,000 bleach-synchronized N2 and GAL257 animals in S-medium supplemented with HB101 *E. coli* bacteria shaking at 20°C and 180 RPM. After four days, gravid adult were bleached in 0.5 M NaOH and 5% NaClO for 5-7 minutes, and released embryos were incubated in M9 buffer overnight. Hatched L1 larvae were fed for 6 hours on NGM plates seeded with HB101 bacteria, after which they were collected and washed five times with M9 prior to processing. Worm dissociation was performed as previously described with minor adjustments^68^. In short, 80 µl of worm pellets were incubated in 500 µl of SDS-DTT solution (0.25% SDS, 200 mM DTT, 3% sucrose and 20 mM HEPES) for four minutes and immediately washed five times with 1 ml of egg buffer (118 mM NaCl, 48 mM KCl, 25 mM HEPES pH 7.3, 2 mM CaCl2 and 2 mM MgCl2). Worm pellets were resuspended in 400 µl of Pronase E (20 mg/ml, Sigma) in egg buffer and incubated for 10 minutes on a roller. Mechanical disruption was carried out by using a 23G needle coupled to a syringe, and pipetting up and down for 10-20 times. Samples were incubated again for 10 minutes on a roller, followed by a mechanical disruption step using the needle and syringe. Digested samples were centrifuged at 100*g* for 1 minute and dissociated cells were collected by transferring the supernatant to a 1.5 ml tube containing 1 ml L15/FBS (10% FBS (Invitrogen) and 1% Penicillin-Streptomycin (Sigma) in L-15 medium (Invitrogen)) to stop the digestion. Worm pellets resulting from the 100*g* centrifugation were resuspended with 400 µl of Pronase E, and additional incubation and mechanical disruption steps were performed, prior to repeating the abovementioned steps to collect the dissociated cells. Cells were then washed once in 1 ml L15/FBS, followed by centrifugation at 1000*g*, 4°C for five minutes, and filtering using a 10 µm strainer (PluriSelect). Filtered cells were washed twice in PBS and fixed with 70% ice-cold ethanol for 1 hour, washed once with a wash buffer 1 (9.5 ml nuclease-free H2O (Ambion), 0.3 ml 5 M NaCl (Ambion), 0.2 ml HEPES (GIBCO), 3.6 µl pure spermidine solution (Sigma-Aldrich), EDTA-free protease inhibitor tablet (Sigma Aldrich), 40 µl 0.5M EDTA (Ambion) and 0.05% Tween-20) with RNASin Plus (1:200, Promega) and frozen in wash buffer 1 supplemented in 10% DMSO and 1:40 RNASin Plus (1:40) for up to 2 months prior using them for RNA-seq and ChIC-seq.

### RNA sequencing

Cells were thawed on ice, washed once in wash buffer 2 (9.5 ml nuclease-free H2O (Ambion), 0.3 ml 5M NaCl (Ambion), 0.2 ml HEPES (GIBCO), 3.6 µl pure spermidine solution (Sigma-Aldrich), EDTA-free protease inhibitor tablet (Sigma Aldrich) and 0.0625% Tween-20) incubated with wash buffer 2 containing Hoechst 34580 (25 mg/ml) for one hour at 4°C on a roller. Cells were washed once in wash buffer 2 prior cell sorting. 500 mCherry-positive or mCherry-negative cells with a 4N Hoechst profile were sorted using a BD FACSAria Fusion into 1.5 ml DNA lo-bind tubes (Eppendorf) containing 5 µl wash buffer 3 (9.5 ml nuclease-free H2O (Ambion), 0.3 ml 5 M NaCl (Ambion), 0.2 ml HEPES (GIBCO), 3.6 µl pure spermidine solution (Sigma-Aldrich), and 0.05% Tween-20). 300 µl of TRIzol reagent (Invitrogen) was added, and samples were stored at -80°C before subsequent processing. RNA extraction, library preparation and sequencing were performed by Single Cell Discoveries B.V. (Utrecht, The Netherlands) using an adapted CEL-seq protocol. Paired-end sequencing was performed in an Illumina NextSeq 500 platform with a depth of 10 million reads per sample. Sequencing reads were aligned to the WBcel235 reference genome (Ensembl release 108) using STARsolo (version 2.7.10a). Analysis of the count matrix was performed in R. Principal component analysis plot was performed using the plotPCA function (DESeq2). Differential gene expression analysis was carried out using DESeq2 package. Lowly expressed genes were filtered out using an average threshold of >10 reads per sample. Differentially expressed genes (DEGs) were defined using a FDR cutoff of 0.05 and log2 fold change cutoff of ± 2. The volcano plot shown in Figure 3C was made using ggplot2. Heatmaps were generated using ComplexHeatmap. The G1/S and G2/M gene lists were made by manually annotating mammalian cell cycle genes, finding *C. elegans* orthologs and classifying them based on their cell-cycle function. The intestinal gene list used was taken from a previously published study^69^. Tissue enrichment analysis was conducted using DEGs as input in Wormbase (https://wormbase.org/tools/enrichment/tea)^70,71^. **Table S2** contains the DESeq log2 fold change and FDR for intestinal versus non-intestinal cells.

### ChIC sequencing

ChIC-seq was performed following a previously described protocol with minor adjustments^42^. Cells were thawed on ice, washed once in wash buffer 1, and incubated overnight in wash buffer 1 containing a primary antibody at 4°C on a roller. The primary antibodies used were against H3K4me3 (1:4000; G.532.8, Invitrogen), H3K9me3 (1:2000; MABI0318, MBL) or H3K27me3 (1:2000; C36B11, CST). Cells were washed once in wash buffer 2, resuspended with wash buffer 2 containing Protein A-MNase (3 ng/ml) and Hoechst 34580 (25 mg/ml) and incubated for 1 hour at 4°C on a roller. Cells were washed twice in wash buffer 2 supplemented with Hoechst 34580 and resuspended with wash buffer 2. 1000 intestinal or non-intestinal cells with a 2N Hoechst profile were sorted into 0.5 ml protein lo-bind tubes (Eppendorf) containing 5 µl of wash buffer 3. The volume containing FACS-sorted cells was adjusted to 10 µl by adding wash buffer 3. The MNase-mediated chromatin digestion was activated by adding 10 µl of wash buffer 3 containing 2 mM CaCl2 and incubating cells for 15 minutes on ice. Digestion was stopped by adding 20 µl of a stop solution (67 µl nuclease-free H2O (Ambion), 8 µl 0.5 M EGTA (Sigma Aldrich), 15 µl IGEPAL CA-630 (Sigma Aldrich) and 10 µl 20 mg/ml Proteinase K (Invitrogen)) and cells were digested for six hours at 65°C followed by heat inactivation at 80°C for 20 minutes and incubation at 4°C until further processing. Blunt-ending, A-tailing and adaptor ligation, *in vitro* transcription and cDNA library preparation steps were performed according to the standard protocol. Equimolar amounts of indexed libraries were sequenced on a NextSeq2000 (Illumina).

Data processing was done using Python following a sortChIC pipeline with minor adjustments. Demultiplexing was performed with demux.py (SMCO v0.1.25), and adaptor trimming was done using cutadapt (version 3.5). Reads were mapped using bwa (version 0.7.17.r1188) and assigned to molecules with bamtagmultiome.py. Files were deduplicated using bamFilter.py and repetitive regions were removed from analysis using bedtools. BigWig files were made using bamCoverage, with a bin size of 20 bp and smooth length of 80 bp. Genomic region windows were exported from IGV. Average plots and heatmaps were generated using computeMatrix and plotHeatmap (deepTools).

### Statistical analyses

Plots of the ZEN-4::GFP levels were generated with ggplot2 using R. Graphs and data analysis of smFISH experiments, intestinal cell and nuclear counts and DAPI quantifications were produced using GraphPad Prism 9 or 10. A minimum of two biological replicates were conducted for each experiment. Statistical significance was defined with unpaired Student *t* test or Mann-Whitney test where appropriate unless otherwise mentioned.

### Data availability

RNA sequencing and ChIC sequencing data are available at Gene Expression Omnibus accession GSE304329 and GSE304344.

## Acknowledgments

We thank members of the Galli and Korswagen groups for helpful discussions; Peter Zeller for help optimizing the ChIC-seq experiments; Anita Semjonova for performing pilot DREAM RNAi experiments; Michela Francesconi for help with the initial ChIC-seq analyses. Anko de Graaf and Pim Tonen from the Hubrecht Imaging Center for support with microscopy; Stefan van der Elst from the Hubrecht FACS Facility for support with cell sorting; Marvin Tanenbaum and Ina Sonnen for critical review of the manuscript. Some strains were provided by the Caenorhabditis Genomics Center, which is funded by NIH Office of Research Infrastructure Programs (P40 OD010440). This work was funded by the European Research Council (ERC) under the European Union’s Horizon 2020 research and innovation program, ERC-Starting grant agreement No. 101040020 and an NWO-Veni fellowship (016.veni.181.016) from the Dutch Research Council (NWO).

## SUPPLEMENTARY FIGURES

**Figure S1:**
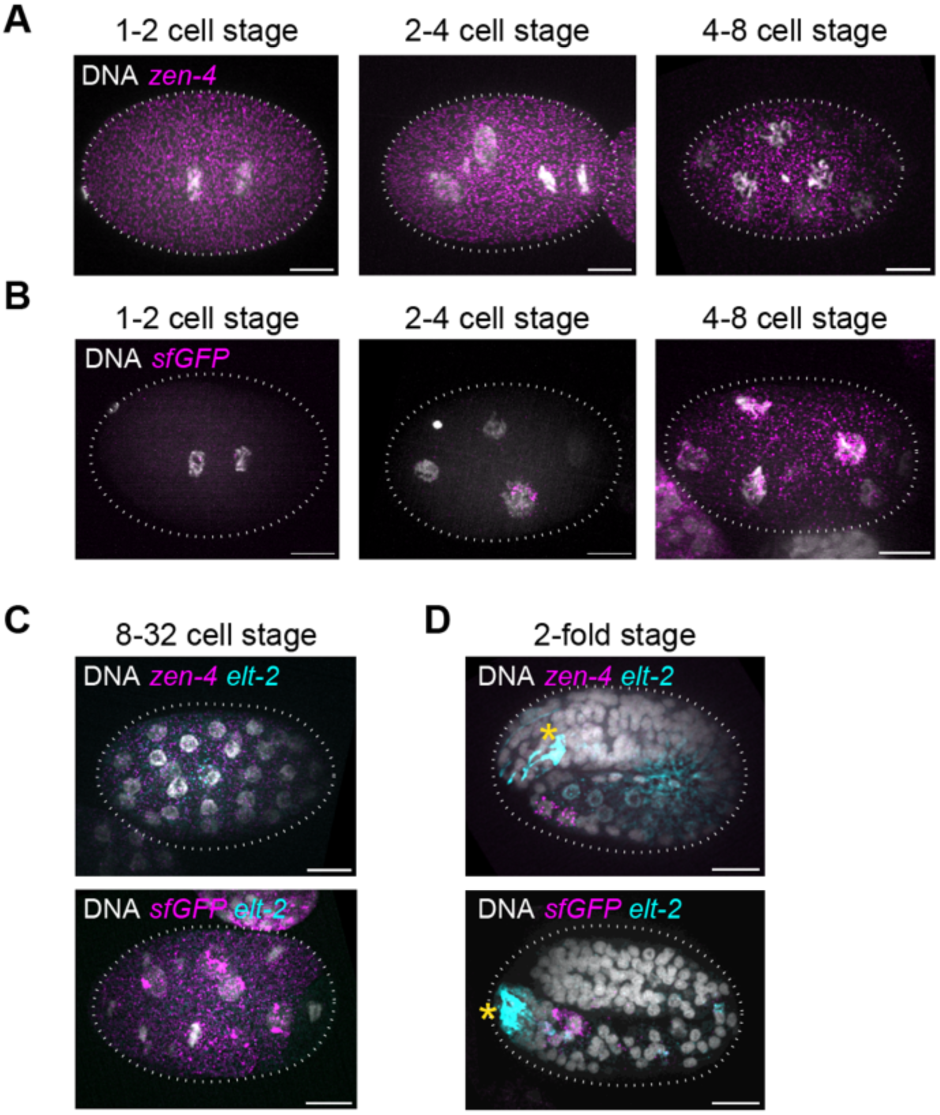
mRNA expression of *zen-4* and *sfGFP* at different stages of embryogenesis. **(A-B)** Representative images of smFISH for *zen-4* **(A)** or *sfGFP* **(B)** mRNA in magenta, and DAPI staining in gray, of early embryos in 1-2, 2-4 or 4-8 cell stages carrying the *zen-4p::^NLS^sfGFP* transgene. **(C-D)** Representative images of smFISH for or *sfGFP* (magenta) and *elt-2* mRNAs (cyan), and DAPI staining (gray) in 8-32 cell stage **(C)** and 2-fold stage embryos **(D)** from a strain carrying the *zen- 4p::^NLS^sfGFP* transgene. Yellow asterisks mark the *lin-48p-TdTomato* co-injection marker. Dashed lines indicate the egg border and scale bars depict 10 μm in all images.

**Figure S2:**
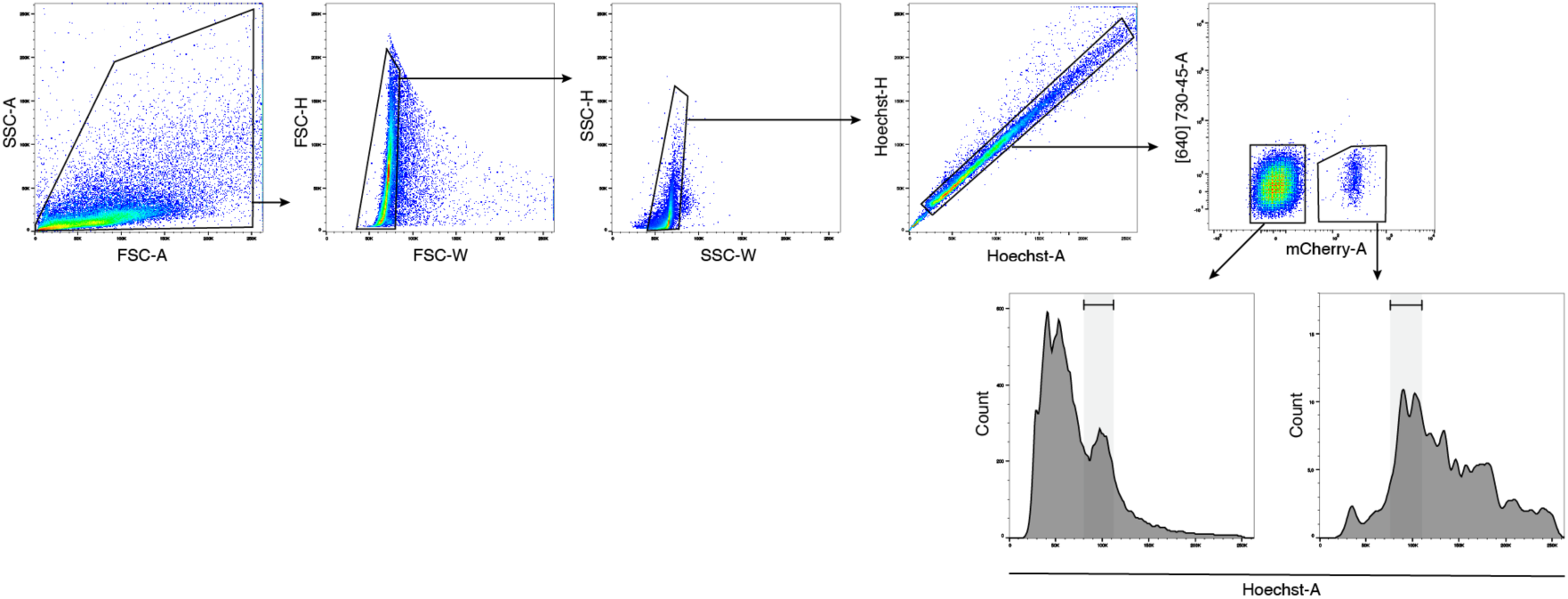
Gating strategy for isolating intestinal and non-intestinal cells in S/G2/M phases.

**Figure S3:**
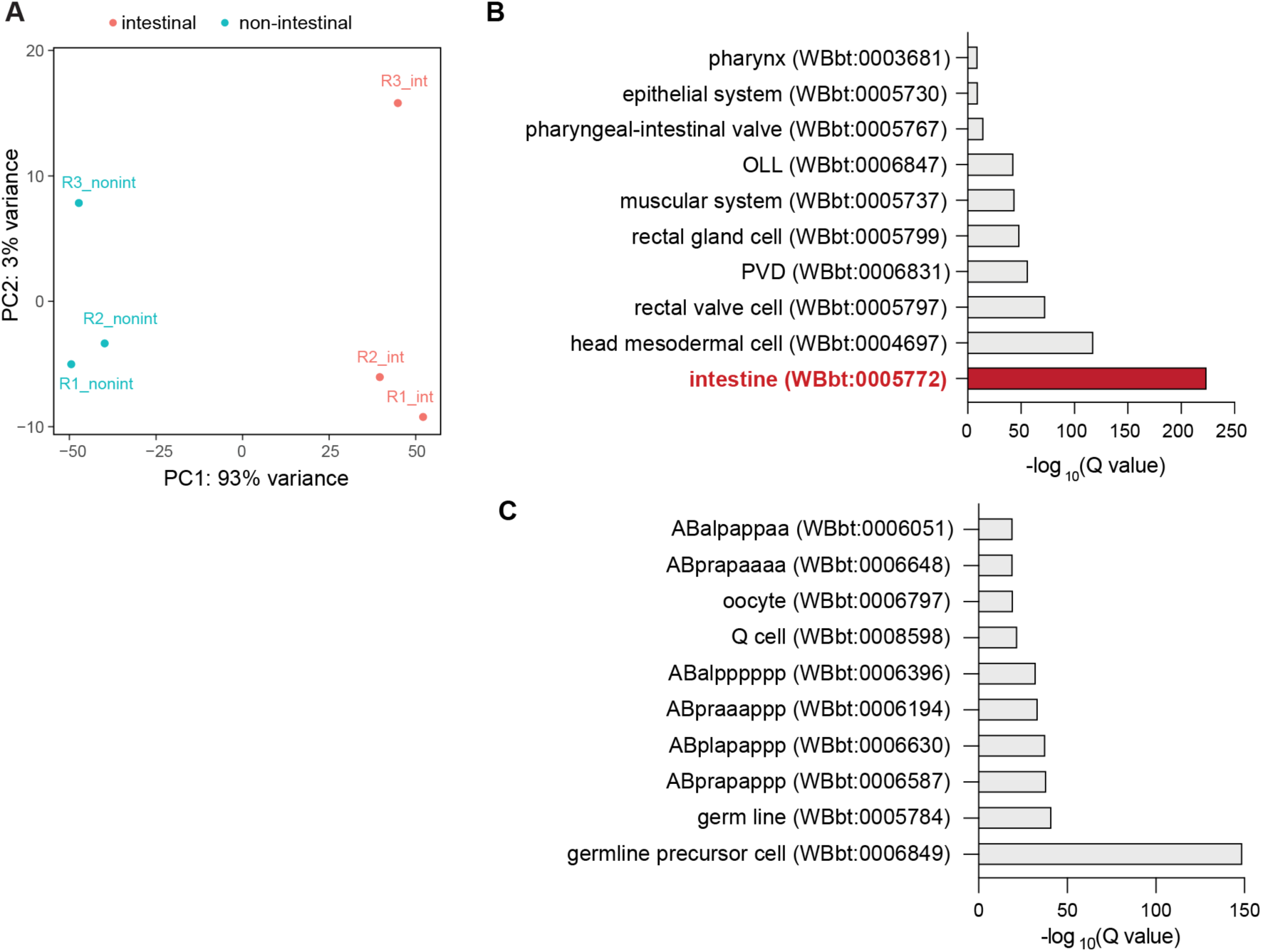
Purified intestinal and non-intestinal cell populations show distinct transcriptional profiles. **(A)** Principal component analysis of the top 500 most expressed genes in individual replicates of intestinal (red) and non-intestinal (blue) cells. **(B-C)** Tissue-enrichment analysis of differentially expressed genes that are significantly enriched in the intestinal **(B)** and non-intestinal **(C)** cell populations showing the 10 most represented tissue terms.

**Figure S4:**
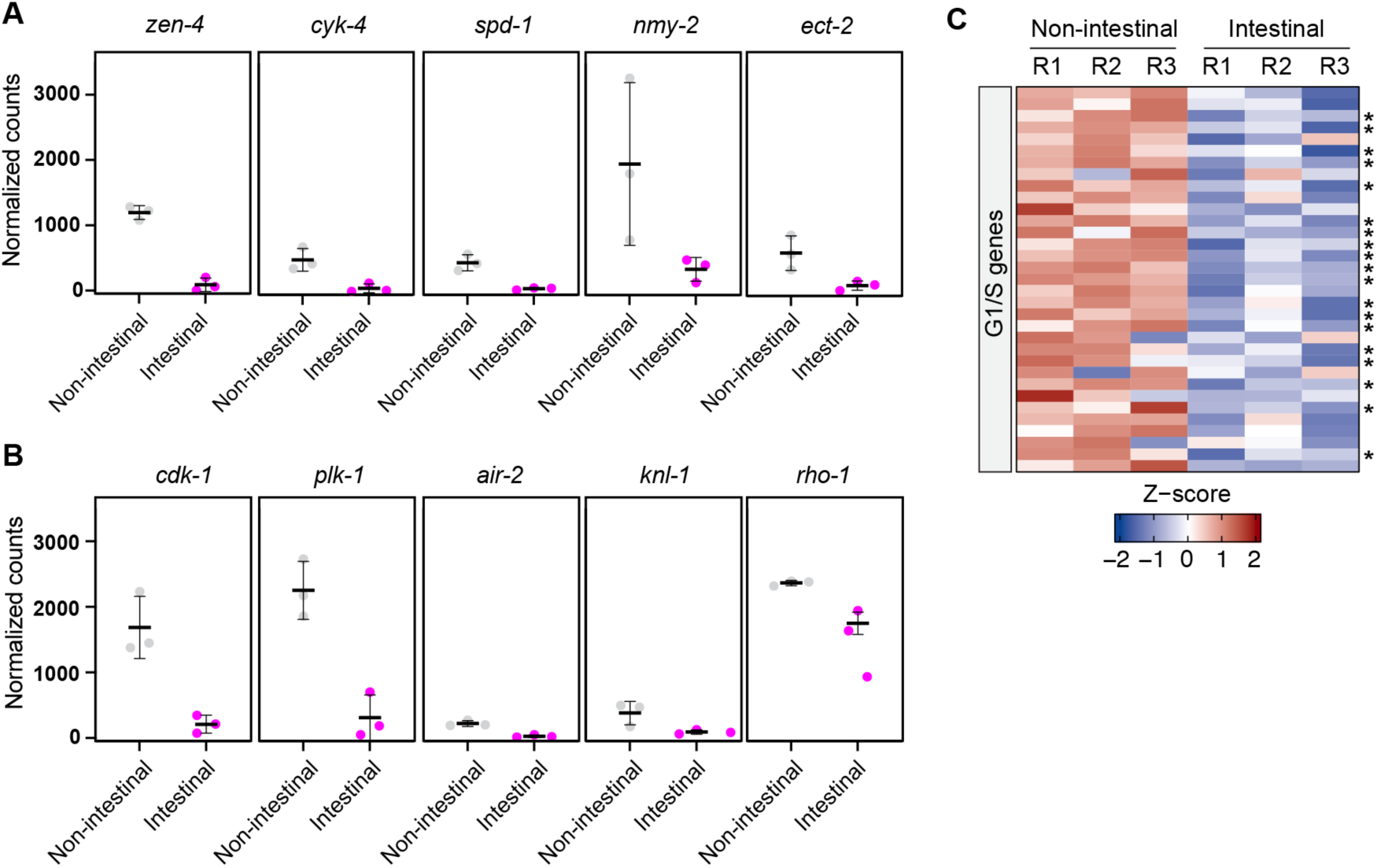
Cell-cycle genes are expressed at lower levels during intestinal endomitosis. (A) Normalized counts of cytokinesis gene transcripts in intestinal and non-intestinal cell populations after DESeq2 normalization. (B) Normalized counts of G2/M transcripts in intestinal and non-intestinal cell populations after DESeq2 normalization. (C) Heatmap depicting the Z-scores and Log2FC of a subset of genes annotated as G1/S genes. Asterisks depict statistical significance.

**Figure S5.**
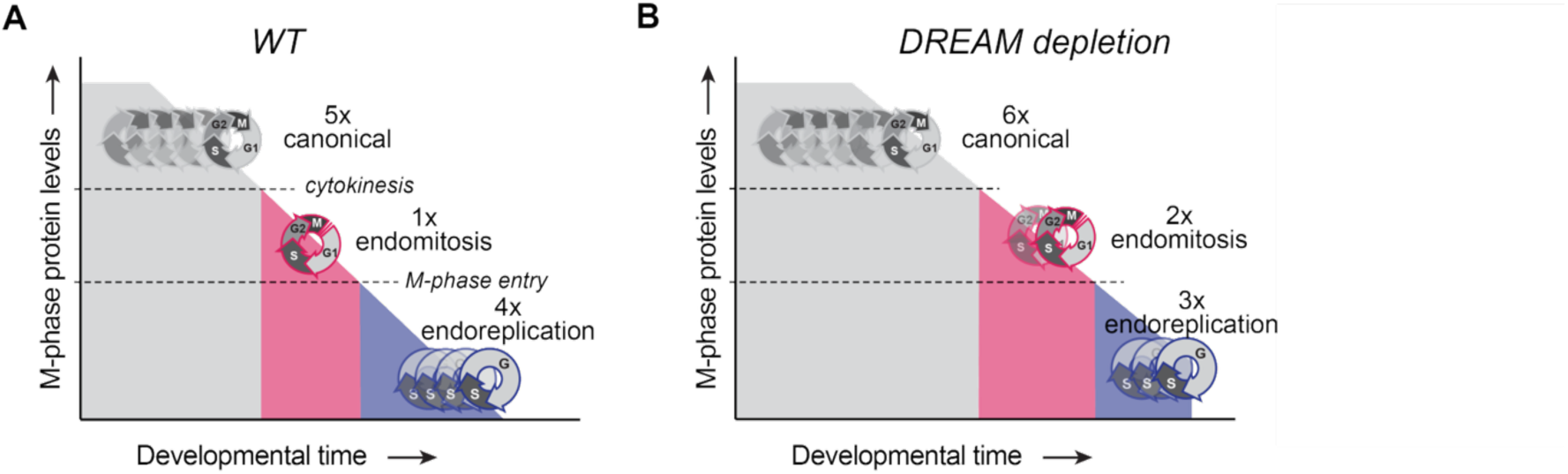
Model for the function of DREAM during *C. elegans* intestinal cell cycles **(A)** In WT animals, the expression of M-phase regulators starts to become repressed at the end of embryogenesis. When the levels of cytokinesis proteins drop below a specific threshold (top dashed line), cells are unable to perform cytokinesis and will perform an endomitosis M phase. When M phase protein levels drop even further (lower dashed line), there will be insufficient M phase factors to enter M phase, and cells will perform endoreplication cycles. **(B)** Upon depletion of DREAM components, the repression of M phase regulators is decreased, leading to sustained levels of M-phase proteins that in turn lead to additional canonical cell cycles during embryogenesis and the conversion of an endoreplication cycle into an endomitosis cycle during larval development.

**Table S1.**
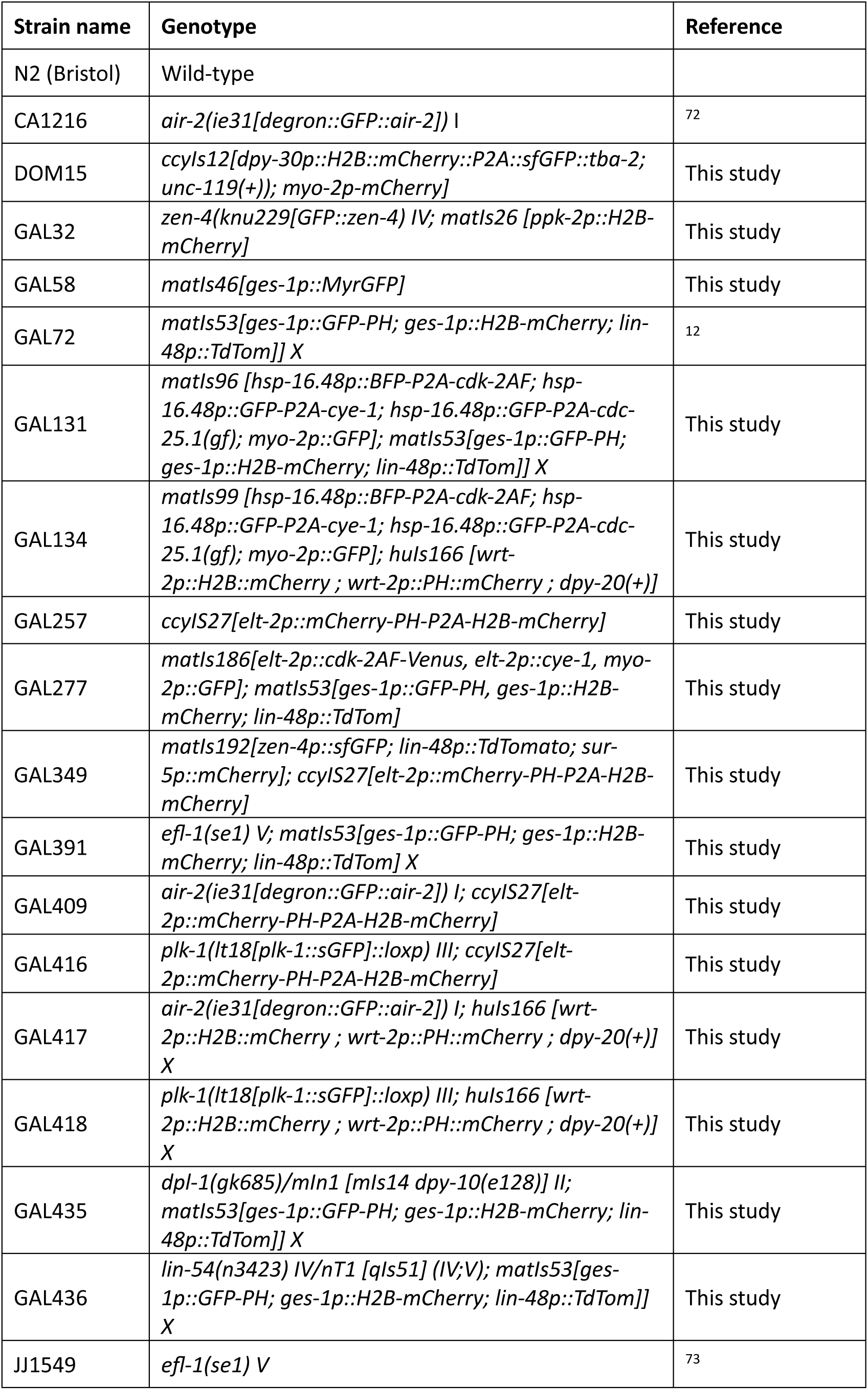
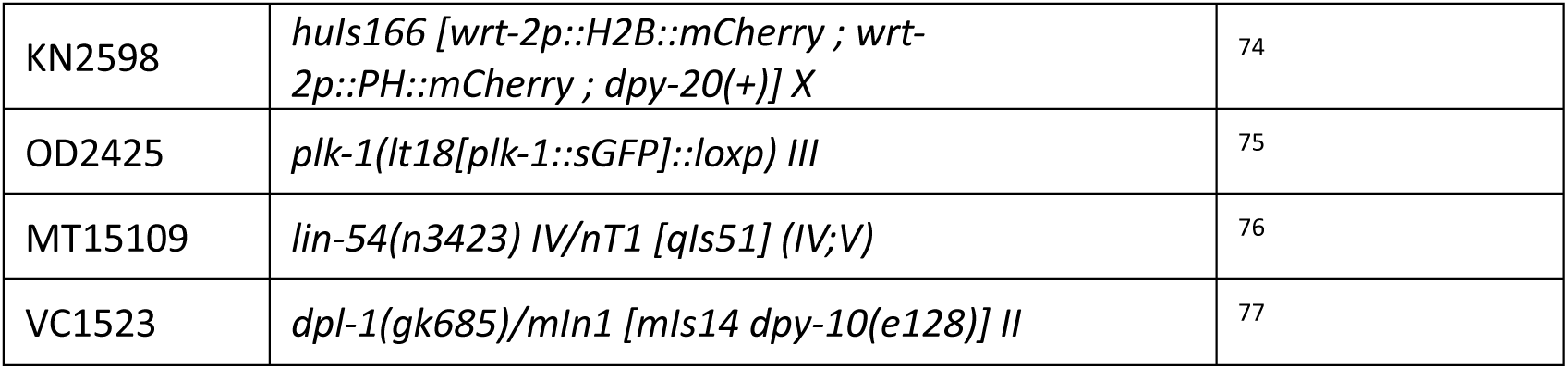
List of strains used in this study.

## Supplementary Files

Table S2

Differential gene expression analysis for intestinal versus non-intestinal cells.

Movie S1

Live imaging of an embryonic intestinal cell undergoing a canonical cell-cycle M phase. Time is relative to metaphase (hh:mm).

Movie S2

Live imaging of an intestinal cell in an L1 larva undergoing endomitosis M phase. Time is relative to metaphase (hh:mm).

## Notes

### Competing Interest Statement

The authors have declared no competing interest.

